# Comprehensive *in silico* analysis reveals candidate regulatory mechanisms underlying selective cerebellar vulnerability in pontocerebellar hypoplasia

**DOI:** 10.64898/2026.09.28.754951

**Authors:** Lízia Branco, Simone Mayer

## Abstract

Pontocerebellar hypoplasia (PCH) is a group of ultrarare, neurodegenerative disorders characterized by cerebellar and pontine hypoplasia. Genetic analysis over the last two decades has revealed an increasing number of pathogenic variants in a wide range of broadly expressed genes functioning in RNA processing, tRNA metabolism, and translation. However, the mechanisms linking these ubiquitous processes to brain region-specific vulnerability are unknown. Here, we established a multi-level variant-to-function *in silico* framework to predict the molecular consequences of PCH-associated variants in TSEN complex genes. These variants were predicted to have heterogeneous effects on diverse protein properties, including stability, subcellular localization, and degradation, supporting variant-specific rather than uniform disease mechanisms. Complementary transcriptomic analyses showed that PCH-associated genes were not globally enriched in the prenatal cerebellum. Instead, their expression was coordinated in a stage- and cell type-specific manner during cerebellar development. We therefore hypothesize that multiple PCH-associated genes are regulated by a common set of transcription factors, providing an explanation of the selective vulnerability of the cerebellum and to the phenotypic convergence of genetically diverse PCH subtypes. In summary, this study prioritizes candidate variants for biochemical, cellular, and in vivo validation, and identifies regulatory programs, cell lineages, and developmental windows for targeted, mechanistically informed disease modelling.

## Introduction

Pontocerebellar hypoplasia (PCH) is a group of rare, autosomal recessive neurodegenerative disorders characterized by a severe hypoplasia of the cerebellum and pons (Van Dijk *et al*, 2018). PCH patients share neuroimaging features, such as the dragonfly appearance of the cerebellum on coronal imaging, as well as motor and cognitive impairments with very limited developmental progress (Namavar *et al*, 2011; Ekert *et al*, 2016). To date, seventeen subtypes of PCH have been characterized (Kukulka *et al*, 2025) and 25 PCH-related genes are currently listed in the OMIM database (Table 1). The known cellular and molecular functions of these genes are heterogeneous but often related to RNA processing or translation (Table 1). It is unknown why defects in these apparently ubiquitous processes result in a brain region-specific pathology and why genes with such distinct functions all converge on similar phenotypic clinical manifestations.

PCH2a (OMIM # 277470) is the most prevalent form of PCH (Kukulka *et al*, 2025), accounting, for example, for an estimated 90% of cases in Germany (PCH-Familie e.V.). It is most commonly caused by a homozygosity of a founder mutation (p.A307S) in *TSEN54* (Budde *et al*, 2008). TSEN54 is a structural subunit of the transfer-RNA (tRNA) splicing endonuclease (TSEN) complex, which catalyses the removal of introns from intron-containing pre-tRNAs (Zhang *et al*, 2023; Trotta *et al*, 2006). The TSEN complex is composed of four protein subunits: two catalytic, TSEN2 and TSEN34, and two structural, TSEN15 and TSEN54. Although less common, pathogenic variants in other TSEN subunits were reported to result in a phenotype similar to PCH2a, granting the subtype names PCH2b (OMIM # 612389), PCH2c (OMIM # 612390) (Budde *et al*, 2008), and PCH2f (OMIM # 617026) (Breuss *et al*, 2016). PCH4 (OMIM # 225753) and PCH5 (OMIM # 610204) are severe TSEN54-associated PCH subtypes typically caused by compound heterozygous combinations of a nonsense or splice site variant and a missense variant in *TSEN54* (Namavar *et al*, 2011).

Animal models of loss of function of *TSEN54* orthologs have so far yielded limited insights into PCH pathogenesis (Kagermeier *et al*, 2024). In zebrafish, morpholino knockdown of tsen54 revealed loss of structural definition in the brain and increased cell death (Kasher *et al*, 2011). In mouse, the complete loss of Tsen54 function is embryonic lethal (Ermakova *et al*, 2018). Drosophila models carrying loss-of-function mutations in the orthologs of *TSEN54* also showed reduced brain lobe volume and increased apoptosis (Schmidt *et al*, 2022). Cross-species modelling of PCH2a is further constrained by the limited evolutionary conservation of the affected position: among these organisms, the residue equivalent to TSEN54 p.Ala307 is only present in mouse (Budde *et al*, 2008).

Despite advances in understanding TSEN complex structure and function, three of the four subunits contain large regions that are unresolved in the available cryo-EM structures and are predicted to be intrinsically disordered (Zhang *et al*, 2023; Yuan *et al*, 2023; Sekulovski *et al*, 2023). Variants mapping to these regions, including the common TSEN54 p.A307S, therefore cannot be interpreted structurally. Biochemical assays showed that pathogenic TSEN variants do not affect the endonuclease activity of the recombinant complex, causing instead a mild thermal destabilization of the heterotetramer (Sekulovski *et al*, 2021). This indicates that impaired tRNA splicing per se may not fully account for disease severity and points to a gap in our understanding of how these variants drive pathology. While the recently developed human cerebellar organoid model of PCH2a (Kagermeier *et al*, 2024) provided first insights into the cellular disease mechanisms, the molecular mechanisms underlying TSEN-related PCH remain poorly understood.

To address this gap, we established an *in silico* framework that systematically predicts the structural and functional consequences of all PCH-associated variants in TSEN complex genes (Fig 1). For each variant, we assessed effects on protein stability, subcellular localization, and the gain or loss of functional sequence motifs, including degrons, eukaryotic linear motifs, and RNA-binding protein motifs (Fig 1). We also compared our *in silico* predictions with the scarce published experimental data, which corroborated some of our *in silico* findings. We found a wide range of predicted changes in all these features depending on the specific variant. Therefore, our protein functional analyses point to heterogeneous, variant-specific consequences rather than a single shared disease mechanism.

**Figure 1.**
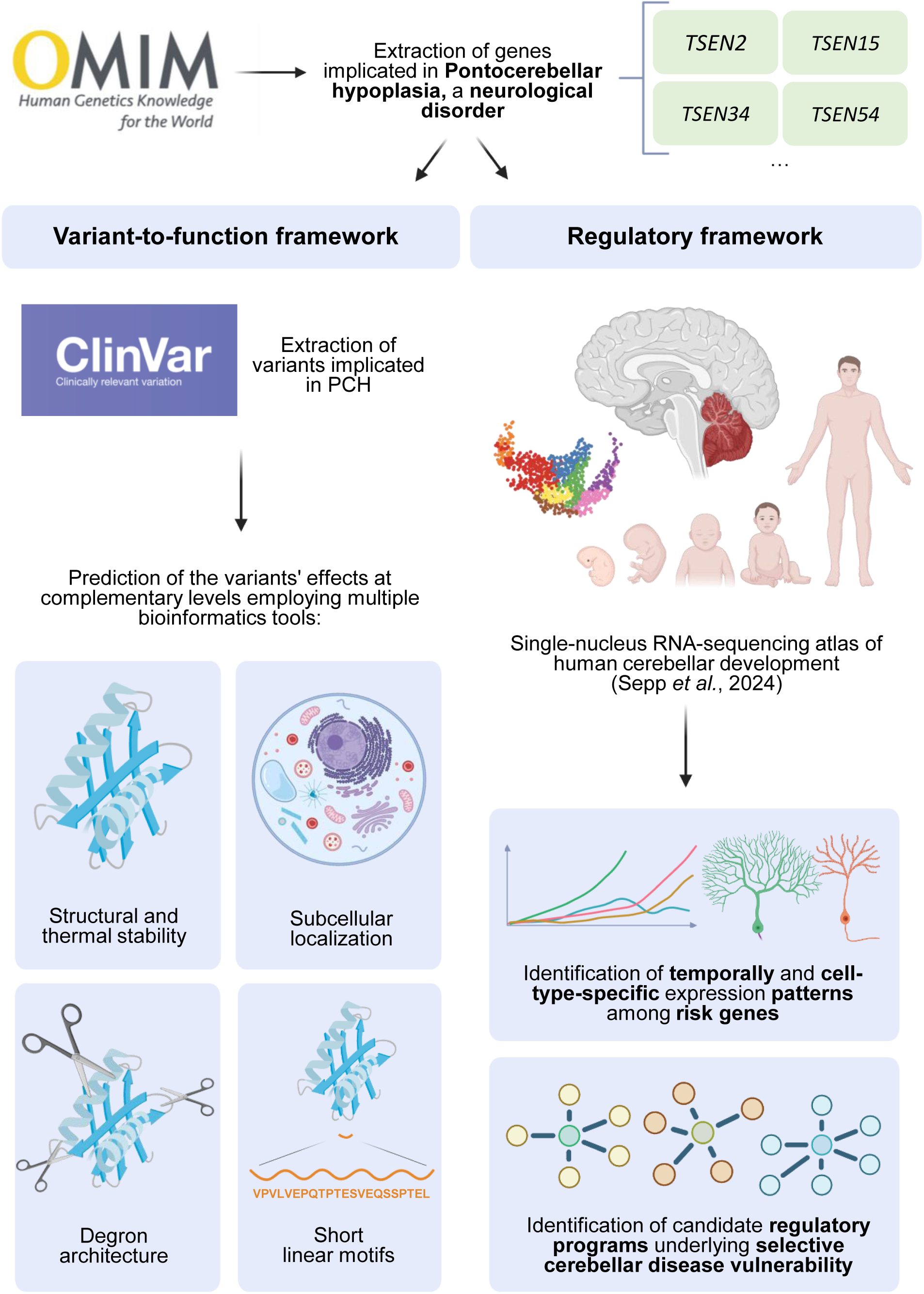
Study overview introducing the multi-level variant-to-function *in silico* framework and the complementary analysis of the spatiotemporal expression of PCH-associated genes during cerebellar development.

To investigate potential mechanisms underlying the specific vulnerability of the cerebellum in PCH, we complemented our variant-centric analyses with global gene expression data analyses. First, given the emerging evidence that TSEN proteins may bind to RNA species beyond tRNA (Hurtig *et al*, 2021; Dhungel & Hopper, 2012; Tsuboi *et al*, 2015), we assessed the RNA-binding propensity of the individual subunits. We identified a previously unrecognized high binding propensity of TSEN34 toward transcripts involved in neurogenesis and neuronal development, such as *ZIC2*, *CBLN1*, and *XKR4*. Second, we leveraged published transcriptomic datasets of human prenatal (Cao *et al*, 2020) and cerebellar development (Sepp *et al*, 2024) to characterize the longitudinal expression and co-expression patterns of the 25 genes currently associated with PCH (Fig 1, Table 1). This analysis revealed 26 candidate transcription factors that we hypothesize to regulate multiple PCH-associated genes in a developmental stage- and cell type-specific manner. Based on this convergent regulation, we propose that the disruption of a single PCH-associated gene may not only impair its direct molecular pathway but also perturb broader cell type-specific regulatory states, thereby propagating the effects of individual mutations beyond their immediate downstream targets, possibly through epigenetic mechanisms. Moreover, our analysis highlighted particularly vulnerable cell types across specific developmental windows – the Purkinje cell lineage prenatally, and the inhibitory neuron, granule cell, and oligodendrocyte lineages postnatally and into adulthood –, offering a possible explanation for the selective hypoplasia.

## Results

### Collection and characterization of ClinVar PCH-linked variants in TSEN genes

ClinVar (Landrum *et al*, 2014) aggregates variant classifications submitted from diagnostic laboratories and expert panels under standardised criteria, making it the most systematic available record of variation in ultra-rare disorders such as PCH. To establish a comprehensive variant dataset for downstream analyses, we therefore extracted all variants in TSEN complex genes (*TSEN2*, *TSEN15*, *TSEN34*, *TSEN54*) annotated in ClinVar to be associated with the PCH subtypes 2a, 2b, 2c, 2f, 4 and 5. This yielded a total of 103 variants (Fig EV1A, B), comprising 81 protein-altering, 6 noncoding or splice-region, and 5 synonymous variants (Fig EV1C). Variants concentrated mostly on the structural subunit *TSEN54*, followed by *TSEN2* (Fig EV1A). Most variants were classified as pathogenic/likely pathogenic and only a minority as benign/likely benign (Fig EV1D).

To accurately interpret disease-associated variants, clinical annotations need to be considered alongside their predicted molecular impact (Richards *et al*, 2015). We therefore evaluated the consistency between ClinVar pathogenicity classifications and those predicted by independent variant effect prediction (VEP) tools. Deleteriousness scores for each variant were retrieved through Ensembl Rest API (Yates *et al*, 2015) and summarised per variant as a weighted average score ranging from 0 (benign-like) through 1 (intermediate) to 2 (pathogenic-like) (Fig EV1E, F; Table 2). For most variants, we found a concordance between computational pathogenicity predictions and ClinVar classifications, although discrepancies remained for a subset of variants (Fig EV1E, F; Table 2).

### Predicted destabilization may contribute to, but does not fully explain, TSEN-related pathogenicity

Alterations in protein stability driven by missense variants are a common pathogenic mechanism in human genetic disease (Stefl *et al*, 2013; Wang & Moult, 2001; Yue *et al*, 2005). We therefore evaluated the impact of all missense variants on protein stability by predicting changes in Gibbs free energy (ΔΔG, kcal/mol) using four complementary widely used computational approaches: FoldX (Delgado *et al*, 2019), mCSM (Pires *et al*, 2014), DynaMut2 (Rodrigues *et al*, 2021), and ThermoMPNN (Dieckhaus *et al*, 2024) (Table 3). Overall, predicted destabilization varied substantially across variants, TSEN subunits, and prediction methods (Table 3). However, most variants in *TSEN54*, *TSEN2*, and *TSEN34* were predicted to have minor effects on protein stability (ΔΔG < 1 kcal/mol) irrespective of the method used, whereas *TSEN15* harboured a comparatively higher proportion of destabilizing variants (Table 3). Notably, the predicted destabilization was not correlated with ClinVar pathogenicity classifications (Fig 2A). These findings suggest that impaired protein stability may contribute to the functional consequences of a subset of variants, and disproportionately so in TSEN15, but is unlikely to represent a universal disease mechanism. For the common TSEN54 p.Ala307Ser variant, for example, we obtained a mean predicted ΔΔG of 0.048 kcal/mol, which is in line with the experimentally determined mild thermal destabilization of the complex in pathogenic variants (Sekulovski *et al*, 2021).

**Figure 2.**
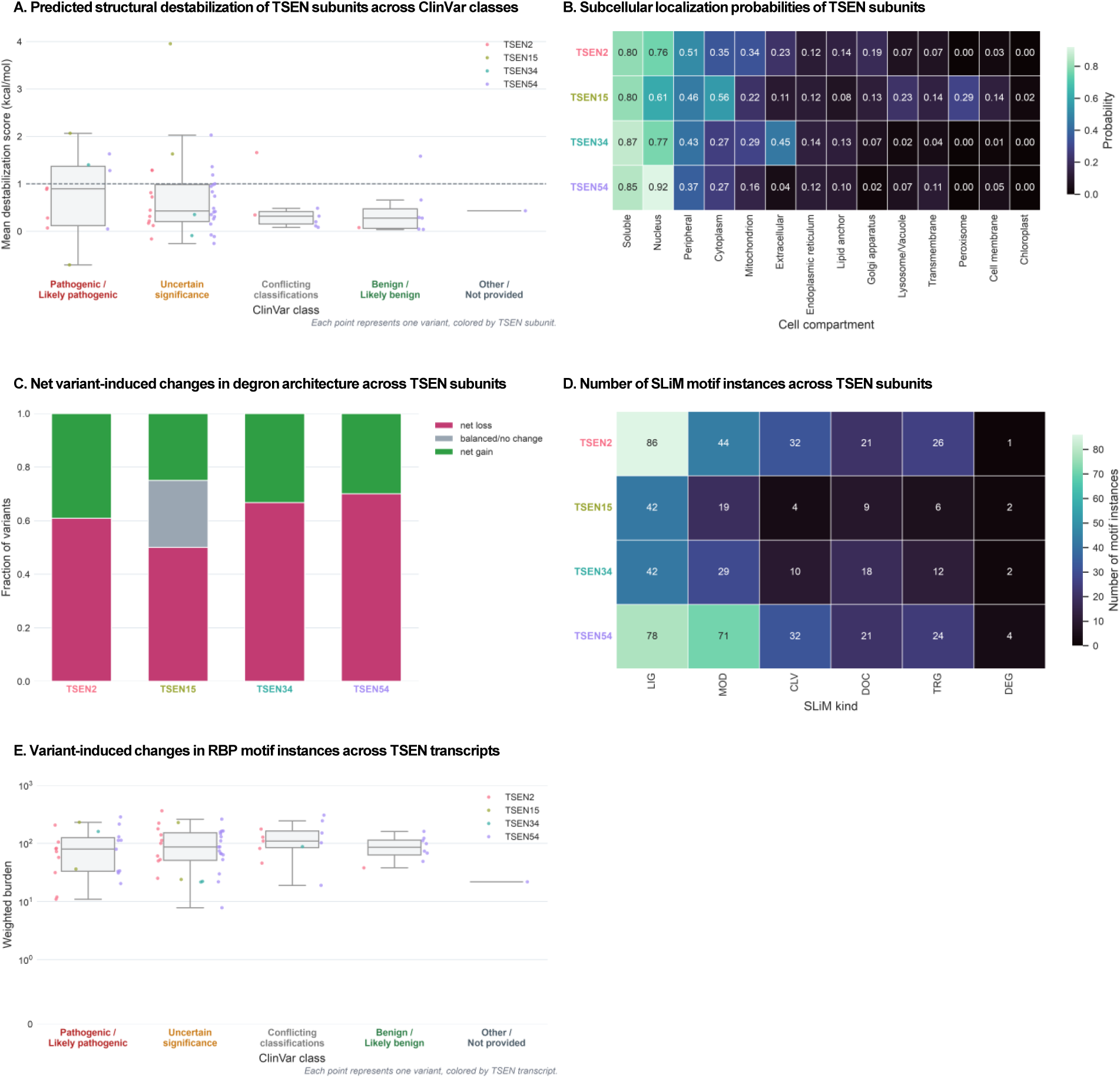
Predicted variant-induced changes in TSEN subunits. A. Predicted structural destabilization of TSEN subunits across ClinVar classes. The prediction score corresponds to the unweighted mean ΔΔG across prediction tools. Each point corresponds to one variant, colored by TSEN subunit; boxes summarize the median and interquartile range within each ClinVar class. B. Subcellular localization probabilities of TSEN subunits, as predicted with DeepLoc2.1. C. Net variant-induced changes in degron architecture across TSEN subunits. Each stacked bar depicts the fraction of variants per TSEN protein classified as net degron loss, balanced/no change (gained degrons exactly equal lost degrons), or net degron gain, based on the difference between mutant and wild-type degron counts, as predicted with Degronopedia. D. Number of short linear motif (SLiM) instances across TSEN subunits, grouped by motif class. E. Variant-induced changes in RNA-binding protein (RBP) motif instances across TSEN transcripts. Weighted burden is the summed absolute-magnitude RBPmap signal shift across all gained, lost, strengthened, and weakened binding-site events. The prediction score corresponds to the unweighted mean ΔΔG across prediction tools. Each point corresponds to one variant, colored by TSEN subunit; boxes summarize the median and interquartile range within each ClinVar class.

### Subcellular localization predictions point to noncanonical TSEN functions

Although TSEN proteins have classically been considered to localize to the nucleus (Melton *et al*, 1980; Lund & Dahlberg, 1998; Paushkin *et al*, 2004), other subcellular localizations have been suggested, such as the cytoplasm and mitochondria (Hayne *et al*, 2023; Dhungel & Hopper, 2012; Yoshihisa *et al*, 2007; Akiyama *et al*, 2022; Takano *et al*, 2005; Zahedi *et al*, 2006). Consistent with this notion, recent large-scale subcellular proteomic profiling identified all four TSEN subunits as predominantly cytoplasmic (Hein *et al*, 2025). To investigate whether altered subcellular localization may contribute to PCH pathogenesis, we used DeepLoc2.1, a tool for predicting the localization probabilities of eukaryotic proteins across cellular compartments (Ødum *et al*, 2024).

All four TSEN proteins were predicted to localize predominantly to the nucleus (61-92%); however, substantial probabilities were also assigned to other cellular compartments (Fig 2B). In particular, all TSEN subunits exhibited a >25% probability of cytoplasmic localization, while all except TSEN54 showed a >20% probability of mitochondrial localization (Fig 2B). In addition, TSEN2 and TSEN34 displayed appreciable probabilities of extracellular localization (>20%), whereas TSEN15 was also predicted to localize to the Golgi apparatus and peroxisomes (>20%) (Fig 2B). We therefore suggest that TSEN subunits may possess functions beyond the canonical tRNA splicing and could potentially act in distinct cellular contexts and functions, in agreement with previously proposed non-nuclear localizations of the complex and/or the respective subunits (Yoshihisa *et al*, 2007; Akiyama *et al*, 2022; Hayne *et al*, 2023; Takano *et al*, 2005; Zahedi *et al*, 2006; Dhungel & Hopper, 2012).

We next assessed whether the pathogenic variants alter the subcellular localization of TSEN subunits. According to DeepLoc2.1’s predictions, subcellular localization was essentially unchanged in most variants, with shifts in compartment probability generally below 1% (Fig EV2, Table 4). Larger shifts (>10%) were confined to truncating variants in TSEN2 and TSEN54, which remove the C-terminal 60-65% of the subunit. Importantly, any altered localization of these truncated proteins depends on whether the corresponding transcripts escape nonsense-mediated decay and are translated. A few missense variants showed intermediate shifts of 5-8%, including TSEN54 p.Lys213Glu and TSEN15 p.Tyr152Cys (Fig EV2, Table 4).

To further investigate mechanisms that could influence protein localization, we employed NLStradamus (Nguyen Ba *et al*, 2009), NLSExplorer (Li *et al*, 2025), and Wregex (Prieto *et al*, 2014) to map nuclear localization signals (NLSs), nuclear import-associated (NIA) regions, and nuclear export signals (NESs) onto each TSEN protein sequence (Fig EV3, Table 4). Variant positions showed minimal overlap with predicted NLS, NIA, and NES motifs (Fig EV3), consistent with the limited localization changes previously predicted by DeepLoc2.1 (Fig EV2, Table 4).

Given that loss of degron sequences is an established mechanism by which variants stabilise their host proteins and cause pathological accumulation (Tokheim *et al*, 2021; Mészáros *et al*, 2017), we additionally mapped degron motifs onto each TSEN protein sequence using Degronopedia (Szulc *et al*, 2024; Prieto *et al*, 2014) (Fig EV3, Table 4). This included both direct degron motifs, i.e., known degron sequences already present within the native protein, and proteolysis-dependent degron motifs, i.e., degrons predicted to emerge at novel N- or C-termini generated through simulated proteolytic cleavage. For each variant we computed net degron balance as the difference between the number of motifs gained and lost (Fig EV4, Table 4). The largest net losses of degron motifs were observed for truncating variants in TSEN2 and TSEN54 (Fig EV4, Table 4). As in the subcellular localisation analysis (Fig EV2, Table 4), however, this net loss likely reflects the extent of sequence loss and should be interpreted in the context of possible nonsense-mediated decay of the respective transcripts. More broadly, predicted changes were smaller but predominantly negative (Fig 2C), ranging between 1 and 26 degron motifs lost (Table 4). PCH-linked variants may therefore generally reduce degradation-promoting sequences in TSEN subunits, potentially stabilizing variant-bearing proteins and the respective fragments released by proteolysis (Tokheim *et al*, 2021; Mészáros *et al*, 2017).

### Possibly altered protein-protein interactions through Short Linear Motifs (SLiMs) driving PCH

Short linear motifs (SLiMs) commonly mediate transient protein-protein interactions and regulatory functions (Davey *et al*, 2011). To assess whether alternations in SLiMs could contribute to altered function of TSEN proteins in PCH, we used the 2024 release of the Eukaryotic Linear Motif resource (Kumar *et al*, 2024) to characterize SLiMs within TSEN proteins and assess the impact of TSEN variants on these features. Six different classes of SLiMs were considered: ligand-binding sites (LIG); post-translational modification sites (MOD); cleavage sites (CLV); docking sites (DOC); targeting signals (TRG); and degradation sites (DEG). TSEN2 and TSEN54 harboured the largest number of predicted SLIMs, most of them of the LIG and MOD motif classes (Fig 1D). Across TSEN subunits, the most abundant predicted motifs were WDR5-binding ligands (required for assembly of H3K4 methyltransferase complexes), Subtilisin/kexin-like proprotein convertase (PCSK) cleavage sites (suggestive of potential regulated proteolytic processing), and ATG8 family interaction motifs (commonly involved in selective autophagy and lysosome-directed vesicular trafficking pathways) (Kumar *et al*, 2024) (Fig EV5A).

We then evaluated the effects of all variants on the SLiMs landscape (Table 5). Predicted motif losses were substantially more frequent than motif gains, with TSEN2 and TSEN54 exhibiting the highest burden of motif alterations (Fig EV5B). As in the degron and subcellular localisation analyses (Fig EV2, EV4; Table 4), however, most disruptions were driven by a small subset of truncating variants in TSEN2 and TSEN54 (Fig EV5C). Among the remaining variants, the most affected SLiM class varied considerably between variants and subunits and net changes in motif instances were correspondingly small, ranging between 0 and 3 motifs lost or gained (Fig EV5C, Table 5). TSEN54 p.A307S, for example, was predicted to gain a CK1 phosphosite and an N-glycosylation site, alongside a positional shift in a USP7-binding motif (Table 5). We therefore suggest that altered protein-protein interactions and post-translational modifications may contribute to PCH pathogenesis, but in a heterogeneous, variant- and subunit-specific manner.

### Altered RNA binding proteins (RBPs) binding sites in TSEN transcripts unlikely to cause pathogenicity

RNAs are bound by proteins that control their maturation, transport, stability, localisation, and translation (Gerstberger *et al*, 2014; Van Nostrand *et al*, 2020) through interactions ranging from sequence-specific recognition to largely non-specific association (Jankowsky & Harris, 2015). Importantly, disruption of these interactions is an established cause of human disease (Lukong *et al*, 2008). To characterize RNA-binding protein (RBP) motif architecture within TSEN transcripts and to determine whether PCH-associated variants tend to disrupt or create such motifs, we used RBPmap (Paz *et al*, 2014). Among the TSEN transcripts, *TSEN34* harboured the largest number of high-confidence RBP-binding sites (Z-score > 3 and P-value < 0.05), whereas *TSEN54* was associated with the lowest number of distinct RBPs (Fig EV6A, Table 6).

We next evaluated the extent to which TSEN variants alter predicted RBP-binding profiles (Table 6). Overall, the cumulative magnitudes of gained and lost signals were broadly comparable across TSEN transcripts (Fig EV6B, Table 6) and variant-induced shifts showed little relation with ClinVar pathogenicity classifications (Fig 1E, Table 6). Collectively, these findings suggest that alterations in predicted RBP binding are not a predominant determinant of variant pathogenicity across TSEN genes.

### Validation of variant-to-function framework across the PCH-associated gene set

To provide an integrated overview of the variant-to-function analyses, we compiled the predicted functional consequences of each variant into a single summary table (Table 7). This consolidated resource enables direct comparison of variant-specific effects across multiple molecular layers and highlights candidate mechanisms that may contribute to TSEN-related PCH pathogenesis for the specific pathogenic variants.

Beyond the TSEN subunits, PCH is genetically heterogeneous and involves genes acting in diverse cellular processes (Van Dijk *et al*, 2018; Kukulka *et al*, 2025). We therefore used three additional PCH-associated genes as a proof-of-concept to demonstrate the broader applicability of our framework. Specifically, we selected *RARS2* (PCH6, Edvardson *et al*, 2007), *EXOSC3* (PCH1B, Wan *et al*, 2012), and *AMPD2* (PCH9, Akizu *et al*, 2013), which, alongside *TSEN54*, are among the genes most frequently reported in association with PCH (Kukulka *et al*, 2025). For each gene, we extracted ClinVar variants classified as pathogenic or likely pathogenic in relation to the corresponding PCH subtype and applied the same variant-to-function workflow (Table 8 and Fig EV7). Concordance with biochemical evidence remains difficult to assess systematically, given the current scarcity of experimental data for these genes. However, one EXOSC3 missense variant, p.Asp132Ala, was recently reported to be thermally destabilizing (Wijeratne *et al*, 2026), consistent with our predicted destabilizing effect for this variant (Table 8).

### TSEN34 preferentially binds mRNAs related to neurodevelopment

The canonical function of the TSEN complex as a tRNA splicing endonuclease requires direct RNA binding. However, emerging evidence suggests that TSEN proteins may participate in additional RNA-associated processes beyond pre-tRNA splicing, including mRNA degradation (Hurtig *et al*, 2021; Tsuboi *et al*, 2015) and rRNA processing (Dhungel & Hopper, 2012). We therefore used catRAPID (Armaos *et al*, 2021) to predict the RNA-binding propensity of TSEN subunits across multiple RNA classes: circular, coding, and small and long non-coding RNAs. Notably, all high-confidence interactions (170 interactions; Z-score > 3) were mediated by TSEN34 (Fig 3A), which presented a particularly high binding propensity to coding RNAs (155 of 170 high-confidence interactions).

**Figure 3.**
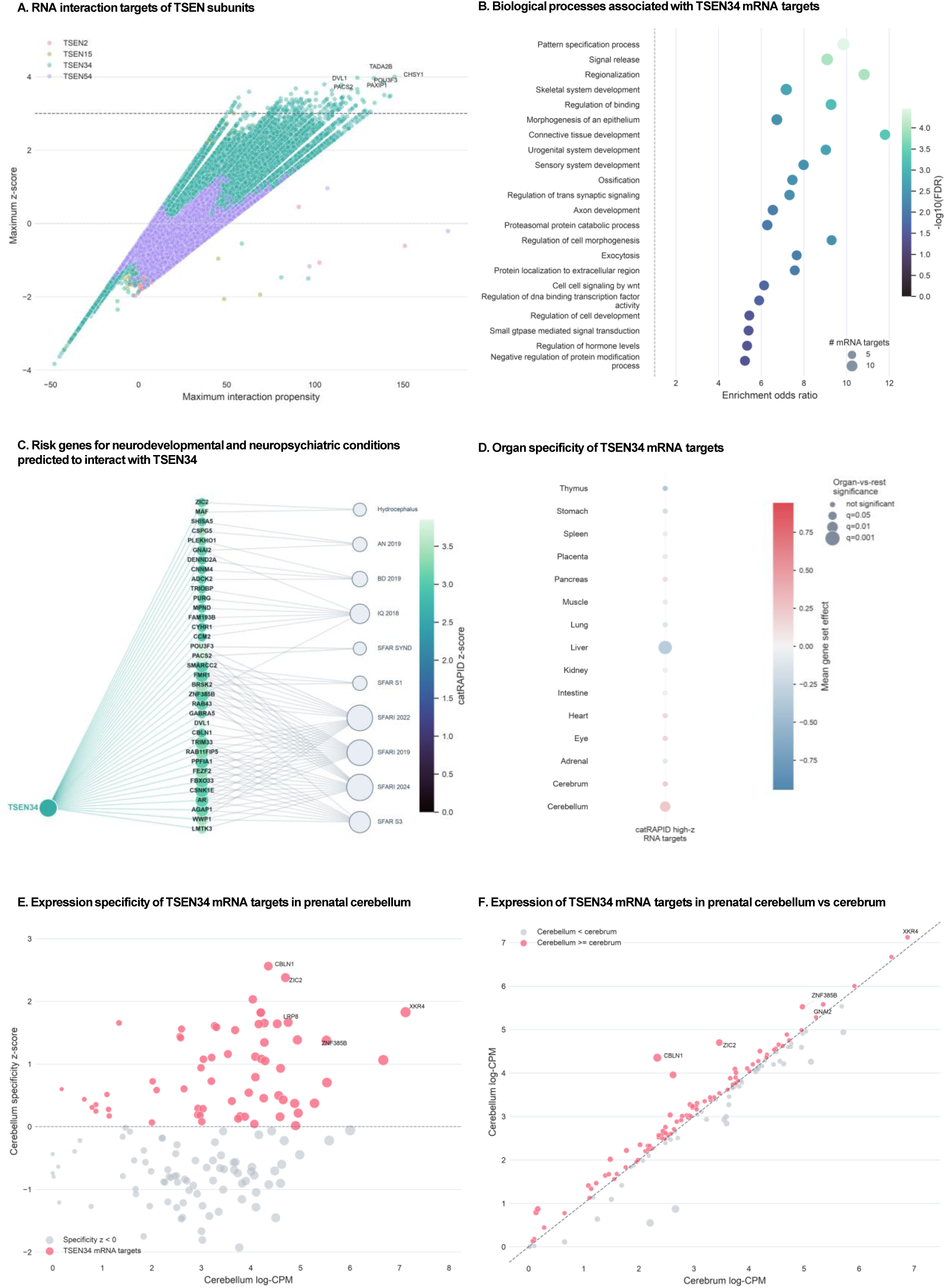
Prediction of TSEN34 binding to transcripts encoding neurodevelopmental genes. A. RNA interaction targets of TSEN subunits. Each point corresponds to the maximum interaction propensity and maximum Z-score obtained for the interaction between a TSEN subunit and an RNA target, as assessed with catRAPID. Points are colored by TSEN subunits and Z-score=3 is marked with a dashed line. The 6 highest-confidence binding partners of TSEN34 are labelled. B. Biological processes associated with TSEN34 mRNA targets. The x-axis shows the enrichment odds ratio and dot color encodes -log_10_ Benjamini-Hochberg FDR from a one-sided hypergeometric test. The displayed terms are the top 20 most enriched biological processes ranked by target-gene overlap, odds ratio, and FDR. C. Risk genes for neurodevelopmental and neuropsychiatric conditions predicted to interact with TSEN34. The considered conditions include SFARI syndromic and S1/S3 gene panels (Abrahams *et al*, 2013) and gene sets for hydrocephalus, anorexia nervosa (AN 2019), bipolar disorder (BD 2019), and IQ (IQ 2018) (Mato-Blanco *et al*, 2025). Gene-node colour encodes maximum catRAPID Z-score (see panel A) and node size reflects the number of matched disorder sets. D. Organ specificity of TSEN34 mRNA targets in a human prenatal transcriptomic atlas (Cao *et al*, 2020). Dot color shows the mean gene-set effect relative to the organ-specific background distribution, with positive values indicating higher expression than expected for that organ and negative values indicating lower expression. Stage-resolved gene-set effects for the displayed organ were compared with all other organs combined using a two-sided Mann-Whitney U test, followed by Benjamini-Hochberg correction across all displayed organ/gene-set combinations. Dot size encodes organ-vs-rest significance as log_10_ Benjamini-Hochberg FDR; non-significant dots are shown at the smallest size. E. Expression specificity of TSEN34 mRNA targets in prenatal cerebellum. The x-axis shows average cerebellar expression as log-CPM and the y-axis shows the gene’s cerebellum specificity z-score relative to its expression across other organs. Genes with negative cerebellum specificity z-scores are shown in grey; genes with non-negative specificity are shown in color. Circle size scales with cerebellar expression. Selected gene labels are ranked across all positive-specificity target genes using a combined score defined as the average of the cerebellum specificity Z-score percentile and the cerebellum log-CPM percentile. F. Expression of TSEN34 mRNA targets in prenatal cerebellum vs cerebrum. The x and y-axis show average cerebellar and cerebrum expression, respectively, as log-CPM. The dashed diagonal marks equal expression in the two organs; points are colored by whether cerebellar expression is lower than cerebrum (grey) or at least as high as cerebrum (red). Circle size scales with the absolute cerebellum-minus-cerebrum difference. Selected gene labels are ranked across all cerebellum-higher target genes using a combined score defined as the average of the cerebellum-minus-cerebrum percentile and the cerebellum log-CPM percentile.

To assess their potential biological function, we performed gene ontology (GO) enrichment analysis on the corresponding gene set. mRNAs predicted to interact with TSEN34 were significantly enriched for biological processes related to development (e.g. pattern specification, regionalization, morphogenesis of epithelium) and neurodevelopmental processes (e.g. regulation of trans synaptic signalling and axon development) (Fig 3B). Additionally, mRNAs related to exocytosis were also enriched (Fig 3B), corroborating the predicted extracellular localization of TSEN34 (Fig 2B).

Interestingly, several predicted target genes (35 of 155 mRNAs predicted to interact with TSEN34) were also risk genes for neurodevelopmental and neuropsychiatric conditions, including hydrocephalus, bipolar disorder and autism spectrum disorder (ASD) (Mato-Blanco *et al*, 2025; Abrahams *et al*, 2013) (Fig 3C). We hence hypothesize that the rare genetic disorder PCH may be mechanistically linked to the more common neurological conditions such as ASD.

We next asked whether this set of high-confidence TSEN34 target RNAs was preferentially expressed in the developing cerebellum. Therefore, we leveraged a published human prenatal single-cell transcriptomic atlas (Cao *et al*, 2020) comprising 4 million cells from 121 samples, 15 organs, and spanning 72-129 postconceptional days. We found this RNAs to be preferentially expressed in both the cerebellum and cerebrum relative to the remaining organs (Fig 3D). A subset of genes was additionally highly expressed and preferentially enriched in the cerebellum (Fig 3E,F). This subset of high-confidence TSEN34 targets included *ZIC2*, *CBLN1*, and *XKR4*, all of which have been implicated in cerebellar development and function. *ZIC2*, together with its paralog *ZIC1*, regulates granule cell proliferation and differentiation and is required for normal foliation of the cerebellum (Aruga *et al*, 2002). *CBLN1* encodes a granule cell-secreted glycoprotein essential for the formation, maintenance, and plasticity of parallel fibre-Purkinje cell synapses (Hirai *et al*, 2005).*XKR4*, a phospholipid scramblase preferentially expressed in the cerebellum, has been associated with variation in cerebellar volume during development (Shook *et al*, 2017).

Therefore, we propose that TSEN34 may preferentially associate with coding transcripts involved in neurodevelopmental and cerebellar-enriched programmes, raising the possibility that its functional repertoire extends beyond its canonical role in tRNA processing. We further hypothesise that such preferential binding to cerebellum-enriched, neurodevelopmentally relevant mRNAs could contribute to the brain region-specific pathology of TSEN-related PCH.

### Cerebellar specificity and developmental expression patterns of PCH-associated genes

PCH is a genetically heterogeneous group of disorders, and mutations in many genes beyond the TSEN subunits, most of which encode proteins involved in broadly expressed cellular processes such as tRNA splicing and mRNA degradation (Table 1), converge on the same characteristic cerebellar phenotype (Van Dijk *et al*, 2018). We therefore asked whether the 25 PCH-associated genes exhibited preferential expression in the developing cerebellum relative to other organs using the human prenatal transcriptomic atlas (Cao *et al*, 2020). At the gene-set level, both TSEN complex genes and the broader set of PCH-associated genes were significantly depleted from the prenatal cerebellum and cerebrum (Fig 4A). Consistent with this, only a small subset of individual PCH-associated genes, namely *PRDM13*, *SLC25A46*, *VPS53*, and *PCLO,* showed positive enrichment in the cerebellum relative to other organs (Fig 4B).

**Figure 4.**
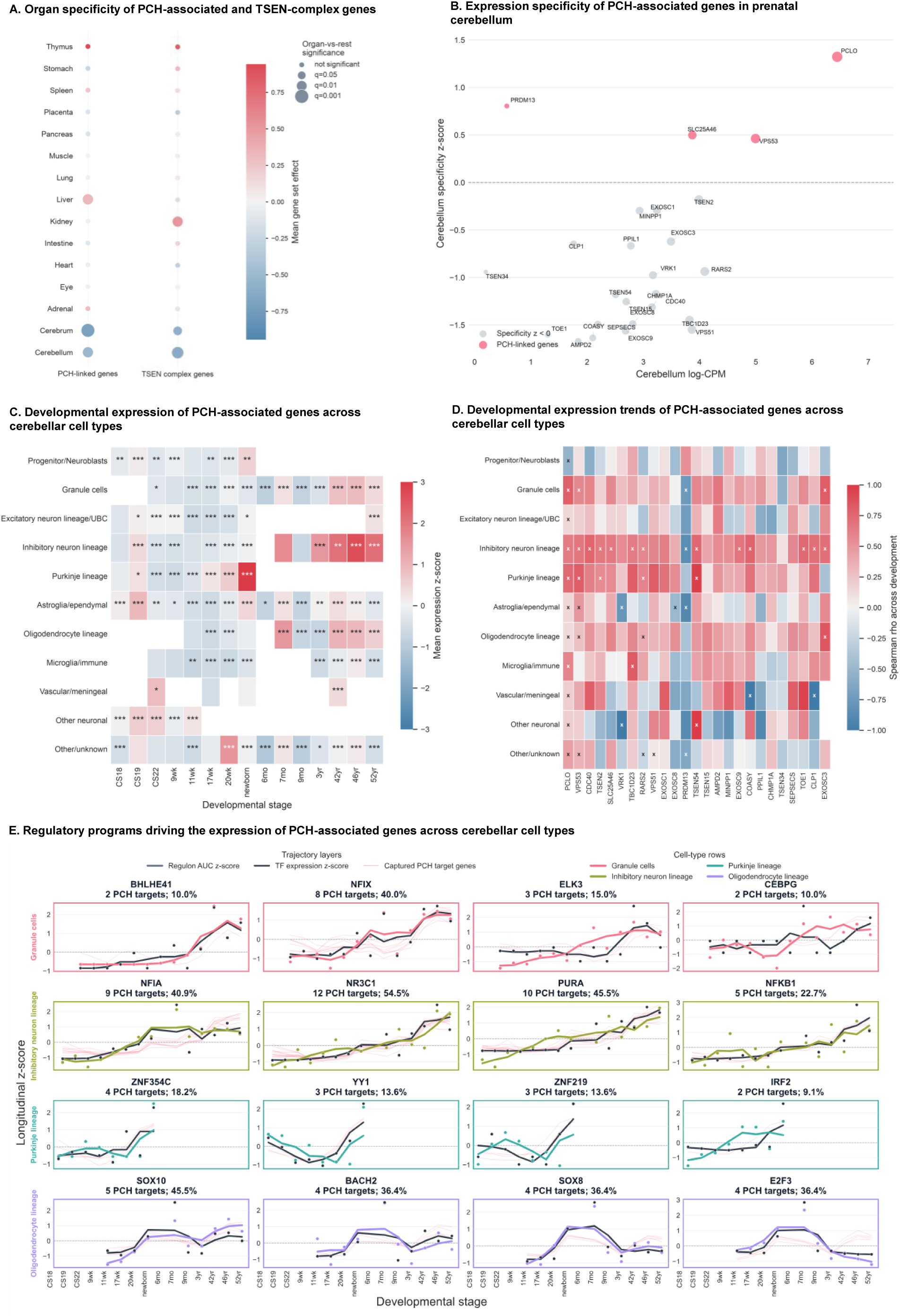
Coordinated, cell-type-specific developmental expression trajectories of PCH-associated genes in the human cerebellum. A. Organ specificity of PCH-associated and TSEN-complex genes in a human prenatal transcriptomic atlas (Cao *et al*, 2020). Dot color shows the mean gene-set effect relative to the organ-specific background distribution, with positive values indicating higher expression than expected for that organ and negative values indicating lower expression. Stage-resolved gene-set effects for the displayed organ were compared with all other organs combined using a two-sided Mann-Whitney U test, followed by Benjamini-Hochberg correction across all displayed organ/gene-set combinations. Dot size encodes organ-vs-rest significance as -log_10_ Benjamini-Hochberg FDR; non-significant dots are shown at the smallest size. B. Expression specificity of PCH-associated genes in prenatal cerebellum. The x-axis shows average cerebellar expression as log-CPM and the y-axis shows the gene’s cerebellum specificity z-score relative to its expression across other organs. Genes with negative cerebellum specificity z-scores are shown in grey; genes with non-negative specificity are shown in color. Circle size scales with cerebellar expression. Gene labels are shown for all 25 PCH-associated genes. C. Developmental expression of PCH-associated genes across cerebellar cell types, as assessed from a single-nucleus transcriptomic atlas of human cerebellar development (Sepp et al, 2024). For each of the 15 considered developmental stages (Carnegie stage 18 through 52 years) and each of the 11 harmonized cell types, the plotted value is the mean expression z-score of the PCH-associated genes. Asterisks indicate cell-type-vs-rest significance within the same developmental stage, assessed by a two-sided Mann-Whitney U test with Benjamini-Hochberg correction across tested cells (* q < 0.05, ** q < 0.01, *** q < 0.001). D. Expression trends of PCH-associated genes across cerebellar cell types. Each cell is coloured by the Spearman correlation (ρ) between ordered developmental stage and mean gene expression for that gene-cell-type pair. Positive values indicate increasing expression across development; negative values indicate decreasing expression. “x” marks pairs with either a significant monotonic trend (|ρ| ≥ 0.55 and Benjamini–Hochberg-adjusted P-value < 0.1, corrected across all pairs) or, in the absence of such a trend, a stage-peaked pattern (expression range across stages ≥ 85^th^ percentile of all pairs). Unmarked pairs met neither criterion. E. Regulatory programs driving the expression of PCH-associated genes across cerebellar cell types. For each panel, the colored thick line depicts the longitudinal Z-score of fixed-regulon AUCell activity; the black line shows the longitudinal Z-score of the TF mRNA expression; and the thin pink lines show longitudinal Z-score trajectories for all PCH-associated genes captured by the corresponding cell-type-specific regulon. All Z-scores are computed within the displayed TF/regulon or gene trajectory across developmental stages; centered three-stage rolling averages are used as visual trend guides, with points marking observed TF/regulon summaries.

We next examined whether PCH-associated genes instead show a stage- or cell-type-specific expression dynamics within the developing cerebellum utilizing a single-nucleus RNA-sequencing dataset spanning human cerebellar development from early neurogenesis to adulthood (Sepp *et al*, 2024). Related cell populations were consolidated into broader categories: progenitors/neuroblasts, granule cells, excitatory lineage/unipolar brush cells, Purkinje cell lineage, inhibitory neuron lineage, astroglial/ependymal cells, oligodendrocyte lineage, microglia, and other neuronal populations (Table 9).

We found expression of PCH-associated genes to peak during the earliest developmental stages (especially Carnegie Stage 19) in most cell types, followed by an overall decline (Fig 4C). From approximately 17 post-conceptional weeks onward, expression increased again in the Purkinje cell lineage, reaching highest levels at the newborn stage (Fig 4C). In contrast, in adulthood, expression was highest in inhibitory neuronal populations, followed by granule cells and oligodendrocyte lineage cells (Fig 4C). Gene-level trajectories showed that these patterns were driven by distinct subsets of PCH-associated genes in each of these cell types (Fig EV7).

Given the apparent coordinated increase in expression across several PCH-associated gene subsets (Fig 4C, Fig EV7), we quantified this trend by correlating mean expression with developmental stage, treated as an ordinal variable, using Spearman’s rank correlation within each cell type (see Methods). This analysis revealed broadly coordinated, generally monotonic expression dynamics among the 25 PCH-associated genes in granule cells, inhibitory neuron lineage cells, Purkinje cell lineage cells, and oligodendrocyte lineage cells (Fig 4D).

Based on the coordinated temporal expression of multiple PCH-associated genes within specific cerebellar developmental windows and cell types, we hypothesized that there may be a shared upstream transcriptional regulation during cerebellar development. Such regulatory programs could potentially contribute to the selective cerebellar vulnerability observed across genetically distinct forms of PCH.

To test this hypothesis, we first inferred gene regulatory networks from the human prenatal transcriptomic atlas (Cao *et al*, 2020) using SCENIC (Aibar *et al*, 2017). We identified TFs that regulate PCH-associated genes and whose regulons display strong cerebellar specificity during early developmental stages, including RFX4, NEUROD2, and SOX10 (Fig EV8A). Several of these regulons also exhibited greater activity in the cerebellum than in the cerebrum (Fig EV8B).

We next asked whether the respective TFs and regulatory programs were specific for certain cerebellar cell types and conserved across a broader developmental timeline. Therefore, we reconstructed gene regulatory networks from the developmental cerebellar dataset (Sepp *et al*, 2024) again using SCENIC (Aibar *et al*, 2017) and focused on cell types exhibiting highly coherent modules of PCH-associated genes: inhibitory neurons, granule cells, and the Purkinje and the oligodendrocyte lineages (Fig 4D, Fig EV7). Candidate TF-cell type pairs were scored based on three criteria: (i) cerebellar specificity of the regulon, (ii) regulation of a coherent module of PCH-associated genes, and (iii) positive longitudinal correlation between TF expression, regulon activity, and target-module expression across development. Applying these criteria identified a restricted set of TF-cell type pairs (Fig 4E, Fig EV8, Table 10). These TF-cell type pairs represent candidate regulatory programs potentially underlying the coordinated expression of PCH-associated genes during cerebellar development. In the inhibitory neuron lineage, for example, the strongest candidates were NR3C1, PURA, and NFIA, which were predicted to regulate 12, 10, and 9 PCH-associated genes, respectively (Fig 4E, Table 10). *NR3C1* encodes the glucocorticoid receptor, a nuclear hormone receptor and ligand-activated transcription factor; *PURA* encodes a highly conserved DNA/RNA-binding protein required for neuronal proliferation, dendritic maturation, and mRNA transport during brain development (Daniel & Johnson, 2018); *NFIA*, together with other Nuclear Factor I (NFI) family members, coordinates neural progenitor differentiation across multiple brain regions, including the cerebellum (Fraser *et al*, 2020). In the granule cell lineage, the strongest candidate was NFIX, predicted to regulate the expression of 8 PCH-associated genes (Fig 4E, Table 10); NFIX, closely related to NFIA, similarly promotes granule neuron precursor differentiation and controls the timing of granule cell maturation during postnatal cerebellar development (Fraser *et al*, 2020).

### Cross-species comparison of developmental expression trends of PCH-associated genes

Human cerebellar development differs substantially from that of other species, in the size and organization of progenitor zones (Haldipur *et al*, 2019), as well as in cell-type composition, maturation dynamics, and gene expression programmes, as defined by recent anatomical and transcriptomic studies (Sepp *et al*, 2024; Akçay *et al*, 2026; Aldinger *et al*, 2021). We have previously suggested that such differences may necessitate detailed disease modelling in human cells, such as brain organoids (Kagermeier *et al*, 2024). We therefore asked whether the coordinated developmental expression patterns observed in the human cerebellum were conserved in other mammalian species.

We leveraged the mouse and the opossum single-nucleus RNA-sequencing datasets from Sepp *et al*, 2024, which span cerebellar development from Theiler Stage (TS) 18 to two months of age in mouse, and from embryonic day (E)14.5 to adulthood in opossum. Compared with the human dataset, cell-type-level differences in PCH-associated gene expression were mild in both species, with the exception of significantly reduced expression of the PCH-associated gene set in mouse Purkinje cells at TS18 (Fig EV9A,B). Similarly, Spearman correlations between developmental stage and expression of PCH-associated genes were weaker overall in both species compared with the human dataset, and tended to be negative (Fig EV9C,D). This indicates that, unlike in human, where most PCH-associated genes showed increasing expression during cerebellar development (Fig 3D, Fig EV8), their developmental trajectories in mouse and opossum were generally flatter or declining. The main exception was mouse Purkinje cells, in which a subset of PCH-associated genes, namely *Pclo*, *Rars2*, *Vps53*, *Tbc1d23*, *Tsen2*, *Tsen15*, and *Tsen54*, showed significant, positive developmental correlations, partially mirroring the trends observed in human (Fig 3D, Fig EV9C). A comparable pattern was not observed in opossum Purkinje cells.

We expect that the species-specific differences in the timing, magnitude, or cellular distribution of PCH-associated gene-expression programs contributes to the difficulty of fully recapitulating the severity and regional selectivity of human PCH phenotypes in animal models.

## Discussion

PCH comprises a genetically heterogeneous group of severe neurodegenerative disorders that converge clinically on hypoplasia of the cerebellum and pons (Van Dijk *et al*, 2018). A central unresolved question in the field is why variants affecting broadly expressed genes, many of which encode proteins involved in fundamental and ubiquitous cellular processes such as RNA processing, tRNA metabolism, and mitochondrial translation, give rise to such a regionally selective pathology (Van Dijk *et al*, 2018). In this study, we addressed this question through two complementary approaches. First, we established an integrative *in silico* framework that combines complementary prediction tools to assess the molecular consequences of PCH-associated variants in TSEN complex genes. Second, we investigated whether PCH-associated genes showed coordinated developmental expression and shared regulatory architecture in the human cerebellum, which could explain the overlapping phenotypes observed across genetically distinct PCH subtypes.

### PCH-associated TSEN variants converge on a heterogeneous, variant-specific consequences at the molecular level

Our variant-to-function analysis indicates that pathogenic and likely-pathogenic variants in TSEN subunits are unlikely to act through a single uniform mechanism. Instead, the predicted consequences were heterogeneous and variant-specific, involving distinct combinations of altered protein stability, subcellular localization, degron architecture, and short linear motif disruption.

Predicted destabilization, for instance, was prominent for a subset of variants but did not account for all pathogenic classifications (Fig 3C), suggesting that impaired protein stability contributes to TSEN-related PCH in a variant-specific manner rather than representing a universal disease mechanism. This interpretation is consistent with the findings of Breuss *et al*, 2016, who showed that three distinct PCH-associated *TSEN15* variants produced convergent disease phenotypes despite differing molecular consequences. One variant did not substantially affect TSEN15 production or stability but appeared to disrupt protein-protein interactions and strongly impair TSEN assembly, whereas the other two variants likely compromised protein folding and stability (Breuss *et al*, 2016). Similarly, Sekulovski *et al*, 2021 demonstrated that recombinant mutant TSEN complexes can still assemble and retain endonuclease activity *in vitro*, while slightly altering the thermal stability of the complex in a variant-dependent manner.

Our degron and SLiM analyses indicate that disease-associated variants may perturb regulatory sequence features involved in protein turnover, proteolysis-dependent fragment handling, ligand binding, cleavage, or post-translational modification processes (Fig 4C,D). These results are particularly relevant because TSEN subunits function as part of a multi-protein complex whose stoichiometry and assembly state appear to be important for efficient pre-tRNA processing (Sekulovski *et al*, 2021). Variants that do not dramatically destabilize an isolated protein may still affect the lifetime, interaction profile, or regulated turnover of the corresponding subunit, thereby impairing TSEN complex function in a cellular context.

Our analysis also suggests that altered subcellular localization is unlikely to represent a general mechanism across all TSEN-related variants. More broadly, however, the predicted localization profiles of wild-type TSEN proteins (Fig 4A), together with recent subcellular proteomic studies (Hein *et al*, 2025) and reports of noncanonical TSEN RNA substrates (Hayne *et al*, 2023), support the possibility that individual TSEN subunits may operate in cellular contexts beyond the canonical nuclear tRNA-splicing reaction, for example cleaving mRNAs that are localized in the cytoplasm or near the mitochondria (Hayne *et al*, 2023).

This scarcity of experimental data validating the predictions by the computational tools underscores the value of a systematic *in silico* framework for generating testable, mechanistically specific hypotheses in genes for which biochemical characterization remains sparse. Ultimately, the pipeline is not intended to replace experimental modelling, but to make such modelling more targeted and mechanistically informed by narrowing the range of plausible variant-level mechanisms and prioritizing candidate variants for biochemical, cellular, or *in vivo* validation

### TSEN34 as a candidate mRNA-binding protein with neurodevelopmental relevance

Among the TSEN subunits, TSEN34 was particularly predicted to interact with coding RNAs (Fig 3A), and the predicted mRNA binding partners were enriched for basic developmental processes, as well as for specific neurodevelopmental processes, such as axon development and trans-synaptic signalling (Fig 3B). Several predicted targets also overlapped with neurodevelopmental and neuropsychiatric disorder risk genes (Fig 3C) and showed preferential enrichment in the developing cerebellum (Fig 3D). These findings are consistent with previous experimental evidence from yeast studies for non-tRNA TSEN substrates, namely mRNAs (Hurtig *et al*, 2021; Tsuboi *et al*, 2015). Future studies should therefore experimentally test whether TSEN34 or other TSEN subunits bind coding RNAs in human neural cells and whether this could be a driver for the brain region-specific pathology.

### PCH-associated genes converge on shared, cell-type-restricted developmental expression programs rather than uniform cerebellar enrichment

By leveraging diverse transcriptomic atlases, we found that PCH-associated genes were expressed in a coordinated, stage- and cell type-specific manner during human cerebellar development (Fig 4E, Table 10). This offers a mechanistic and testable explanation both for the cerebellar vulnerability of PCH and for the overlapping phenotypes observed across PCH subtypes (Van Dijk *et al*, 2018). Because the proposed convergence operates at the level of transcriptional regulation, single-cell chromatin accessibility profiling (Buenrostro *et al*, 2015) would establish whether these candidate transcription factors occupy accessible regulatory elements at PCH-associated gene loci, and whether accessibility at those elements is altered in disease. Interestingly, epigenetic disturbance has been shown to play a key role in neurodevelopmental disorders, such as Rett syndrome, caused by variants in the methyl-CpG-binding protein MeCP2 (Li *et al*, 2020), and a form of autism spectrum disorder caused by variants in a chromatin remodeler CHD8 (Shi *et al*, 2023). Importantly, comparable effects can arise from genes with no direct role in chromatin regulation. In vanishing white matter disease, recessive variants in any of the five subunits of the translation initiation factor eIF2B (Leegwater *et al*, 2001; van der Knaap *et al*, 2002) reduce eIF2B activity and activate the integrated stress response, driving sustained expression of the ATF4-regulated transcriptional programme (Abbink *et al*, 2019; Wong *et al*, 2019). Although eIF2B is required in every cell, pathology is confined to the white matter and primarily affects astrocytes (Dooves *et al*, 2016), which has been attributed in part to a subset of astrocytes displaying constitutive ISR activity that is further amplified by eIF2B mutations (Wong *et al*, 2019).

### Purkinje cells emerge as a convergent point of vulnerability in early cerebellar development, while later-stage vulnerability extends to inhibitory neurons, granule cells, and oligodendrocyte lineages via candidate regulatory programs

Our regulatory network analysis identified Purkinje cells as the most vulnerable cell type during prenatal cerebellar development (Fig 4C, Fig EV10C). Coordinated developmental expression trends among PCH-associated genes were observed in Purkinje cells in both human and mouse cerebellum (Fig 4D, Fig EV10C) making them a potentially informative cell type for studying early PCH onset across human and mouse model systems. Given the central role of Purkinje cells in organizing cerebellar circuitry and influencing granule cell expansion and maturation (van der Heijden & Sillitoe, 2021), early Purkinje-cell dysfunction could also contribute secondarily to later abnormalities in other cerebellar populations. At later developmental stages, our analyses point to inhibitory neuron lineage cells, granule cells, and oligodendrocyte lineage cells as additional candidate vulnerable populations (Fig 4C, Fig EV8). This suggests that later-emerging disease-relevant processes may involve inhibitory circuit development, granule-cell expansion and maturation, or myelination-associated programs, depending on the affected gene and the regulatory program in which it participates. For example, *TSEN54*, the gene most frequently associated with PCH2, was included in multiple candidate regulons, including NFIA and PURA-associated regulons in the inhibitory neuron lineage and KLF9-associated regulon in granule cells and the Purkinje lineage (Table 9). By contrast, *RARS2* and *CLP1*, associated with PCH6 and PCH10, respectively, were linked to candidate regulons in the inhibitory neuron and oligodendrocyte lineages (Table 9). Our hypothesis thus aligns with neuropathological studies reporting variable loss of Purkinje cells and internal granule cells in PCH2 (Joseph *et al*, 2014; Barth *et al*, 2007, 2), as well as abnormal or delayed myelination in other PCH subtypes, namely PCH6 and PCH10 (Boczonadi *et al*, 2014; Ngoh *et al*, 2016; Karaca *et al*, 2014). On the other hand, the identification of *NFIA* and *PURA*-associated regulons is particularly noteworthy because both genes have been independently implicated in human neurodevelopmental diseases (Afshar *et al*, 2026; Senaratne & Quintero-Rivera, 1993). Heterozygous pathogenic variants in *PURA* cause *PURA*-related neurodevelopmental disorder, a severe developmental encephalopathy characterized by neonatal hypotonia, developmental delay, seizures, and impaired neuronal development (Afshar *et al*, 2026). Similarly, *NFIA* haploinsufficiency causes a neurodevelopmental syndrome characterized by corpus callosum abnormalities, ventriculomegaly or hydrocephalus, developmental delay, hypotonia, and seizures (Senaratne & Quintero-Rivera, 1993). While these predictions do not establish direct causality, they support the hypothesis that PCH vulnerability may involve disruption of shared transcriptional networks governed by neurodevelopmentally relevant TFs, rather than isolated dysfunction of individual disease genes.

### Cross-species comparisons suggest human-specific amplification of PCH vulnerability

Our cross-species comparison also has implications for disease modelling. The observation that PCH-associated genes show weaker group-wide coordination and milder cell-type-specific enrichment during opossum and early mouse cerebellar development (Fig EV10) suggests that some aspects of PCH vulnerability may be more pronounced in the human developmental context. This interpretation is supported by recent studies highlighting features that are unique to human cerebellar development, particularly the relative abundance and gene expression programmes of specific cell types (Becker *et al*, 2026; Aldinger *et al*, 2021). Notably, the developing human cerebellum has been shown to harbour transient, human-specific progenitor populations and developmental programmes that are not observed in mouse (Haldipur *et al*, 2019), and a comprehensive cell atlas of human cerebellar development has revealed cytoarchitecturally distinct regions and developmentally transient cell types that diverge from those described in the mouse cerebellum (Aldinger *et al*, 2021). More specifically, human Purkinje cell neurogenesis has been shown to occur within an outer subventricular zone that is absent from the mouse cerebellum, and to proceed over a markedly compact developmental window (Haldipur *et al*, 2019). Together, these species-specific differences in cerebellar cell-type composition and maturation timing support the possibility that PCH-associated developmental vulnerability may be more pronounced or differently timed in the human cerebellum than in other species, underscoring the value of complementary human-based models, such as cerebellar organoids (Runnebohm *et al*, 2025; Kagermeier *et al*, 2024) and primary prenatal transcriptomic references.

### A two-level, molecular-to-systems model of PCH pathogenesis

Overall, our findings support a two-level model of PCH pathogenesis. At the molecular variant level, disease-associated mutations likely impair protein stability, localization, turnover-associated motifs, and regulatory SLiMs, in a gene- and variant-specific manner. Interestingly, many of these genes are essential for mammalian development (e.g. the Tsen54-/-mouse is embryonic lethal, Ermakova et al., 2019), and we therefore propose that there is a subtle loss of function associated with the pathogenic variants. Interestingly, altered binding of the TSEN complex to proteins through SLiMs and to coding RNAs, represent promising hypotheses for follow-up experimental studies. At the developmental systems level, genetically distinct PCH-associated genes may converge on shared cerebellar transcriptional programs, creating windows of heightened vulnerability in specific cell types. Cell-type-resolved perturbation models should therefore be established experimentally to determine whether the candidate TF-regulon programs identified here (Table 10) mediate this shared vulnerability across PCH subtypes. In this context, the recurrent WDR5-binding motifs predicted across TSEN subunits is of interest, as WDR5 is involved in epigenetic remodelling (Bryan *et al*, 2020) and could therefore provide a mechanistic link between the molecular and developmental systems level understanding of pathogenesis. Together, our two-level, molecular-to-systems model reconciles the molecular heterogeneity of PCH with its convergent clinical phenotype and provides a basis for prioritizing future experimental validation.

### Limitations of this study

First, the predicted effects of PCH-associated variants through diverse *in silico* tools require experimental validation in relevant biochemical, cellular, and developmental model systems. Although the integration of multiple computational tools provides a systematic framework for variant prioritization, these predictions should be interpreted as hypothesis-generating.

Second, each bioinformatic tool used in this study has intrinsic limitations. For example, ELM-based short linear motif prediction depends on the scope and annotation depth of the underlying motif database (Kumar *et al*, 2024). Therefore, additional SLiMs may remain undetected. Similarly, predictions of protein stability, subcellular localization, degron architecture, and RNA-binding propensity are model-dependent and may be influenced by cellular state and tissue-specific regulatory mechanisms that are not modelled in the underlying algorithms.

Third, our analyses primarily evaluated variant effects at the level of individual TSEN proteins or transcripts. We did not model how variants alter the structure, assembly, dynamics, or catalytic function of the complete TSEN complex. This was partly limited by the presence of poorly resolved or intrinsically disordered regions within TSEN subunits (Sekulovski *et al*, 2023; Hayne *et al*, 2023; Yuan *et al*, 2023), which complicates reliable complex-level structural modelling. As a result, variants with limited predicted effects at the individual protein level may still impair TSEN complex assembly, substrate recognition, catalytic activity, or interactions with additional protein partners that were not modelled here.

## Methods

### General computational analysis setup

Core pipeline scripts were run with Python 3.10.7. Main Python packages included pandas 2.2.3, NumPy 1.23.2, SciPy 1.10.1, matplotlib 3.9.2, seaborn 0.13.2, networkx 2.8.6, requests 2.28.1, and statsmodels 0.14.0. R analyses were run with R 4.4.3, using Seurat 5.3.0, SeuratObject 5.2.0, Matrix 1.7.3, dplyr 1.1.4, tidyr 1.3.1, readr 2.1.5, stringr 1.5.1, ggplot2 3.5.2, forcats 1.0.0, AnnotationDbi 1.66.0, org.Hs.eg.db 3.19.1, clusterProfiler 4.12.6, UCell 2.8.0, SingleCellExperiment 1.26.0, data.table 1.18.2.1, and patchwork 1.3.2. Gene regulatory network inference was performed with pySCENIC 0.12.1 in a dedicated Python 3.12.3 environment. Tools accessed through public web-services were run using default settings unless when stated. The respective web-service endpoints and locally cached web outputs were reviewed for the last time on 24 June 2026.

### Variant cohort definition and annotation

ClinVar (Landrum *et al*, 2014) (https://www.ncbi.nlm.nih.gov/clinvar/) records were queried and curated to define a cohort of variants in TSEN-complex genes associated with pontocerebellar hypoplasia (PCH). The TSEN variant cohort was restricted to ClinVar condition labels specifically matching the PCH subtypes relevant to TSEN-complex disease: PCH2a, PCH2b, PCH2c, PCH2f, PCH4, and PCH5. Broad or ambiguous labels, including generic pontocerebellar hypoplasia terms not resolvable to these subtypes, were excluded unless the TSEN gene context allowed unambiguous assignment of a broad PCH2 label to the appropriate OMIM subtype. For each retained variant, all matching PCH subtype labels and condition terms were preserved. ClinVar clinical significance labels were normalized the following harmonized classes: pathogenic/likely pathogenic, uncertain significance, conflicting classifications, benign/likely benign, and other/not provided. When multiple PCH-relevant ClinVar classifications were available for the same molecular variant, the variant was represented once and assigned the most pathogenic retained classification according to a predefined severity ranking. Variants were classified by molecular consequence, including missense, synonymous, nonsense/stop-gained, frameshift, splice-region or intronic, deletion, and other protein-altering or non-protein-altering categories where applicable.

Variants that could not be mapped unambiguously to the relevant coding, transcript, or protein sequence were retained in the variant-level metadata but excluded from analyses requiring a valid mutant sequence. For exploratory comparison with non-TSEN PCH genes, the same framework was applied separately to pathogenic or likely pathogenic ClinVar variants in *RARS2*, *EXOSC3*, and *AMPD2*, corresponding to PCH6, PCH1B, and PCH9, respectively. PCH-associated genes used in developmental and regulatory analyses were curated from the OMIM phenotypic series PS607596 for pontocerebellar hypoplasia and linked to their corresponding PCH subtype.

### Protein and transcript sequences

Human wild-type protein FASTA sequences for TSEN2, TSEN15, TSEN34, and TSEN54 were retrieved from UniProtKB/Swiss-Prot (https://www.uniprot.org/). Ensembl REST (https://www.ensembl.org/) was used to retrieve gene metadata, canonical transcript identifiers, and genomic locus FASTA sequences. Mutant protein sequences were generated computationally when the protein HGVS annotation could be applied unambiguously to the wild-type UniProt sequence. For RNA-level analyses, RefSeq transcript accessions present in ClinVar variant names were used to retrieve transcript records, and mutant transcript sequences were reconstructed when the coding HGVS representation and transcript context allowed unambiguous sequence editing. The same sequence-generation logic was used for the isolated *RARS2*, *EXOSC3*, and *AMPD2* test pipeline where applicable.

### Variant-effect prediction

Variant deleteriousness was evaluated using annotations obtained through the Ensembl VEP REST API (https://rest.ensembl.org). The VEP REST requests included canonical transcript, HGVS, AlphaMissense, CADD, REVEL, conservation, LOEUF, and dbNSFP-style fields when available. The resulting table retained source-scale values for SIFT, PolyPhen-2, REVEL, AlphaMissense, CADD, phyloP, phastCons, GERP++, gnomAD allele-frequency fields, and the LOEUF gene-level constraint metric. These annotations were harmonized at the variant level. For tools returning categorical interpretations, predictions were mapped to benign/tolerated, intermediate/uncertain, or deleterious/pathogenic classes. For continuous predictors, tool-specific thresholds were used to assign an equivalent ordinal class. Each available interpretable call was encoded as 0 for benign, 1 for intermediate or uncertain, and 2 for deleterious/pathogenic. For each variant, the average predictor score was calculated across predictors with available interpretable results; missing predictors were excluded from the denominator and reported separately.

### Structural stability prediction

Structural stability analyses were restricted to variants compatible with single-residue missense modelling, requiring a valid wild-type residue, mutant residue, protein position, and structural template. The structural templates used were retrieved through the AlphaFold DB (Cheng *et al*, 2023) model version 6 API (https://alphafold.ebi.ac.uk/) using UniProt accessions and downloaded as PDB files from the corresponding AlphaFold DB file URLs. Stability effects were predicted with multiple complementary methods: FoldX 4 (Schymkowitz *et al*, 2005) using local RepairPDB and BuildModel workflows, mCSM (Pires *et al*, 2014) through the public mCSM stability-prediction batch endpoint (https://biosig.lab.uq.edu.au/mcsm/stability_prediction_list), DynaMut2 (Rodrigues *et al*, 2021) through its public prediction API (https://biosig.lab.uq.edu.au/dynamut2/api/prediction_list), and ThermoMPNN (Dieckhaus *et al*, 2024) using a local installation of the Kuhlman Lab ThermoMPNN inference workflow. For consistency across tools, predicted changes in Gibbs free energy (ΔΔG) or tool-specific stability outputs were converted into a shared destabilization orientation where positive values indicate predicted destabilization and negative values indicate predicted stabilization. For each variant, predictor-specific ΔΔG or destabilization values were retained, and consensus summaries were calculated across available tools. Structural consensus was evaluated using a 1 kcal/mol destabilization threshold. Variants with at least two scored tools were classified as consistently low-destabilization, consistently destabilizing, or mixed depending on whether all available tools fell below or above the threshold or produced discordant calls. These summaries were stratified by TSEN protein and compared with ClinVar pathogenicity classes.

### Subcellular localization, transport signals, and degron analysis

Subcellular localization was predicted using the DeepLoc 2.1 (Ødum *et al*, 2024) web service hosted by DTU Health Tech (https://services.healthtech.dtu.dk/services/DeepLoc-2.1/). Wild-type and mutant protein FASTA sequences were submitted in batches through the service’s webface CGI endpoint as a multipart file upload, with the “Fast” model and “short” output format. For each protein, wild-type localization probabilities were summarized across predicted cellular compartments. For each mutant sequence, localization changes were calculated as mutant probability minus wild-type probability for the corresponding protein and compartment, expressed in percentage points.

Transport-signal annotations were obtained using complementary NLS/NES approaches. Nuclear localization signals were predicted with NLStradamus (Nguyen Ba *et al*, 2009) (http://www.moseslab.csb.utoronto.ca/NLStradamus/), run locally on the wild-type protein sequence alone, and with NLSExplorer (Li *et al*, 2025) (http://www.csbio.sjtu.edu.cn/bioinf/NLSExplorer/), which was submitted the wild-type protein sequence together with its corresponding AlphaFold monomer model. Nuclear export signal candidates were annotated using a local implementation of the published Wregex (Prieto *et al*, 2014) NES consensus regex pattern (https://ehubio.ehu.eus/wregex/home.xhtml); because the public Wregex web interface was insufficiently stable for automated batch submission, its original PSSM-based scoring was not reproduced, and candidate NES regions reported here reflect consensus-pattern matches only. Predicted transport-signal intervals were mapped onto each protein together with variant positions. Degron annotations were obtained with Degronopedia (Szulc *et al*, 2024; Prieto *et al*, 2014) (https://degronopedia.com/), including annotation of proteolysis-dependent degron regions where available. For each mutation, degron gain and loss were determined by comparing predicted degron intervals in the mutant sequence against the corresponding wild-type sequence. A net degron balance was calculated as the number of gained degron features minus the number of lost degron features, such that negative values indicate predominant degron loss and positive values indicate predominant degron gain.

### Short linear motif analysis

Short linear motifs (SLiMs) were annotated using ELM motif definitions downloaded from http://elm.eu.org/ (Dinkel *et al*, 2012) and scanned locally against wild-type and mutant protein sequences. Wild-type motif architecture was first characterized for each TSEN protein by counting predicted motif instances and grouping them by SLiM functional kind and feature name. Variant-induced motif changes were then quantified by comparing motif instances between the wild-type and matched mutant sequence, matched by identical SLiM kind at the same sequence interval. Motifs present in the wild type but absent after mutation were classified as lost, whereas motifs absent from the wild type but present in the mutant were classified as gained. This coordinate-based matching does not reconcile a motif that shifts position without changing content; the variant cohort reaching this analysis step comprises only missense and nonsense substitutions, which preserve or truncate residue numbering without repositioning retained motifs, so this limitation does not affect the results presented here. Net SLiM balance was calculated as gained motifs minus lost motifs, both globally and by SLiM kind.

### RNA-binding protein motif analysis

RNA-binding protein motifs were assessed with RBPmap (Paz *et al*, 2014) (https://rbpmap.technion.ac.il/) using wild-type and mutant transcript sequences. RBPmap was run with the Human/Mouse motif catalogue, and wild-type TSEN transcripts were first characterized by the number of high-confidence RBP motifs, the number of distinct RBPs represented, and the overall site burden. Mutation-associated effects were then evaluated by comparing RBPmap motif calls between matched wild-type and mutant transcript sequences. Mutation impact was quantified using a continuous weighted signal designed to capture four classes of change: gained motifs, lost motifs, retained motifs with increased signal, and retained motifs with decreased signal. For each variant, gained and lost binding sites were weighted according to Z-score × −log_10_(P-value), whereas retained sites were weighted based on ΔZ-score multiplied by the average −log_10_(P-value) across the WT an_d_ variant sequence. Net directional summaries classified each variant as dominated by increased/gained RBP signal, dominated by decreased/lost RBP signal, or mixed when neither direction accounted for at least 25% of the total weighted signal. Reported motif positions correspond to RNA/transcript coordinates.

### catRAPID RNA-interaction analysis

Potential RNA interaction partners of TSEN subunits were evaluated using catRAPID omics v2.1 protein-versus-transcriptome predictions (Armaos *et al*, 2021) (https://tools.tartaglialab.com/new_submission/catrapid_omicsv2_protein). TSEN proteins were queried against Homo sapiens RNA libraries representing coding RNAs, long non-coding RNAs, small non-coding RNAs, and circular RNAs. RNA targets with catRAPID interaction Z-score greater than 3 for at least one TSEN protein were retained as high-confidence predicted RNA interactors, and for each target gene the strongest TSEN-associated Z-score was recorded. Of the four TSEN proteins queried, only TSEN34 yielded RNA targets meeting this threshold; TSEN2, TSEN15, and TSEN54 had no qualifying high-z interactors, so all downstream GO enrichment, network, and organ-specificity analyses in this section are specific to TSEN34. Gene Ontology Biological Process (GO-BP) enrichment among high-Z target genes was assessed using a custom over-representation analysis. The foreground consisted of 170 high-Z target genes, and the background consisted of all 49,066 genes detected in the corresponding catRAPID output. GO-BP gene sets were obtained from the MSigDB v7.5.1 c5.go.bp collection, retaining only terms containing 5 to 500 background genes. For each GO-BP term, enrichment was tested using a hypergeometric test, and P-values were adjusted using the Benjamini-Hochberg false-discovery-rate procedure. High-Z RNA targets were also overlapped with curated neurodevelopmental disorder risk-gene sets to identify predicted TSEN RNA partners previously implicated in neurodevelopmental and neuropsychiatric conditions (Mato-Blanco *et al*, 2025; Abrahams *et al*, 2013).

### Cross-organ and developmental single-cell expression analysis

Cross-organ prenatal expression specificity was evaluated using in a previously published human prenatal transcriptomic atlas (Cao *et al*, 2020). Gene expression was summarized as average log-normalized expression per organ. For each gene, cerebellar specificity was calculated by comparing cerebellar expression against the distribution of expression across other organs, yielding a cerebellar specificity Z-score. Cerebellum-versus-cerebrum expression differences were calculated from average expression estimates in the two tissues. These analyses were performed separately for TSEN-complex genes, OMIM-curated PCH-associated genes, and catRAPID high-Z RNA target genes. For gene-set organ-specificity plots, stage-resolved gene-set effects for a displayed organ were compared with all other organs combined using two-sided Mann-Whitney U tests, followed by Benjamini-Hochberg correction.

Developmental cerebellar expression was analysed using single-cell transcriptomic atlas of human, mouse, and opossum cerebellar development (Sepp *et al*, 2024). Data from 44-year-old adult donor was excluded from the human dataset because, unlike the neighbouring 42- and 46-year-old samples (14,102 and 6,214 granule cells, respectively, out of 15,580 and 8,033 total cells), it contained zero granule cells and an atypical overall cell-type composition. Original cell-type annotations were mapped to harmonized cerebellar lineages (Table 9), including granule cells, Purkinje lineage, inhibitory neuron lineage, excitatory neuron lineage/UBC, progenitor/neuroblast populations, oligodendrocyte lineage, astroglial/ependymal cells, microglia/immune cells, vascular/meningeal cells, and other categories where retained. Stage-cell-type groups with fewer than 20 cells were excluded from stage-cell-type expression summaries. In the opossum dataset, PCH-associated human gene symbols were mapped to opossum orthologs via Ensembl BioMart human-to-opossum orthology queries, preferring one-to-one orthologs with the highest reported orthology confidence. This yielded 22 of 25 mappable PCH-associated genes. The three unmapped genes (*COASY*, *EXOSC8*, *TSEN15*) had a valid ortholog call whose Ensembl gene ID was not present among the features of this dataset build.

For each gene, mean expression was calculated within each retained stage-cell-type group and converted to a gene-wise Z-score across all retained groups. Heatmap values for PCH-associated genes represent the mean of these gene-wise expression Z-scores across the target set in each stage-cell-type context. Statistical testing compared expression scores for a given target set in one cell type against all other cell types from the same developmental stage using two-sided Wilcoxon/Mann-Whitney tests, followed by Benjamini-Hochberg correction. Significance was denoted as * q < 0.05, ** q < 0.01, and *** q < 0.001.

Longitudinal gene-expression trends were quantified using Spearman correlations (ρ) between ordered developmental stage and mean gene expression within each harmonized cell type, computed independently for each dataset (human, mouse, opossum). A gene-cell-type pair was classified as showing a meaningful trend if either of two criteria was met: (i) a monotonic trend, defined as a Benjamini-Hochberg-adjusted Spearman p-value (q) below 0.1 together with ρ at least 0.55 in magnitude (increasing for ρ ≥ 0.55, decreasing for ρ ≤ -0.55), with the Benjamini-Hochberg correction applied jointly across all gene-cell-type pairs within that dataset; or, failing that, (ii) a non-monotonic but strongly stage-peaked pattern, defined as an expression range (maximum minus minimum mean expression across stages) at or above the 85^th^ percentile of ranges observed across all gene-cell-type pairs in that same dataset. All other pairs were classified as not showing a strong monotonic trend. Trajectory plots display observed stage-level mean expression values with centered three-stage rolling averages used only as visual guides.

### SCENIC-based regulatory network analysis

Gene regulatory networks (GRNs) were inferred with pySCENIC (Aibar *et al*, 2017) 0.12.1 using the human hg38 transcription factor list, motif annotations motifs-v10nr_clust-nr.hgnc-m0.001-o0.0.tbl, and hg38 motif-ranking databases hg38_10kbp_up_10kbp_down_full_tx_v10_clust and hg38_500bp_up_100bp_down_full_tx_v10_clust. Our analysis considered two complementary SCENIC layers. First, an all-organ SCENIC/AUCell context was generated to quantify regulon activity across prenatal organs and to estimate cerebellar specificity using the human prenatal transcriptomic atlas (Cao *et al*, 2020). Cerebellar specificity of the respective transcription factors (TFs) was evaluated both relative to all other organs and specifically relative to cerebrum. Second, the previously employed single-cell transcriptomic atlas of human cerebellar development (Sepp *et al*, 2024) was analysed using one fixed all-cells regulon/AUCell matrix for comparable developmental activity trajectories, together with broad-cell-type GRNs to define cell-type-specific transcription factor-to-target edges.

Candidate hubs were ranked by an integrated score combining five components with fixed weights: the prenatal cerebellum regulon specificity score (floored at zero; weight 4), the percentage of the coherent PCH-associated module captured by the regulon in that cell type (weight 0.04), a combined developmental trajectory-support term consisting of the mean of the regulon-AUCell-versus-module and TF-expression-versus-module correlations (each floored at zero; weight 2), and the statistical significance of the coherent-module overlap test itself, expressed as -log_10_ of its Benjamini-Hochberg-adjusted P-value (weight 0.15). For visualization of multi-layer candidate evidence, a curated subset of these metrics was converted to column-wise Z-scores.

## Data availability

All original analysis code produced in this study will be made publicly available on GitHub upon publication.

## Author contributions

L.B. conceptualization, methodology, formal analysis, data curation, writing - original draft; S.M.: conceptualization, funding acquisition, resources, supervision, writing - review & editing.

## Acknowledgements

We would like to thank Daria Andreeva, Isabel Potthof, Dr. Theresa Kagermeier, Dr. Francesco Castagnetti, and Dr. Magdalena Stock for their valuable feedback on the manuscript. The authors used Claude (Opus 5; Anthropic) to improve the language and readability of the text. All content was subsequently reviewed and edited by the authors, who take full responsibility for the content of the publication.

The research was funded by the German Research Foundation (Deutsche Forschungsgemeinschaft, DFG) under Germany’s Excellence Strategy 2082/2 390761711.“ Icons in figures 1 and expanded view 10 were created in BioRender: (https://BioRender.com/3nvr3b1).

## Figures

**Figure Expanded View 1.**
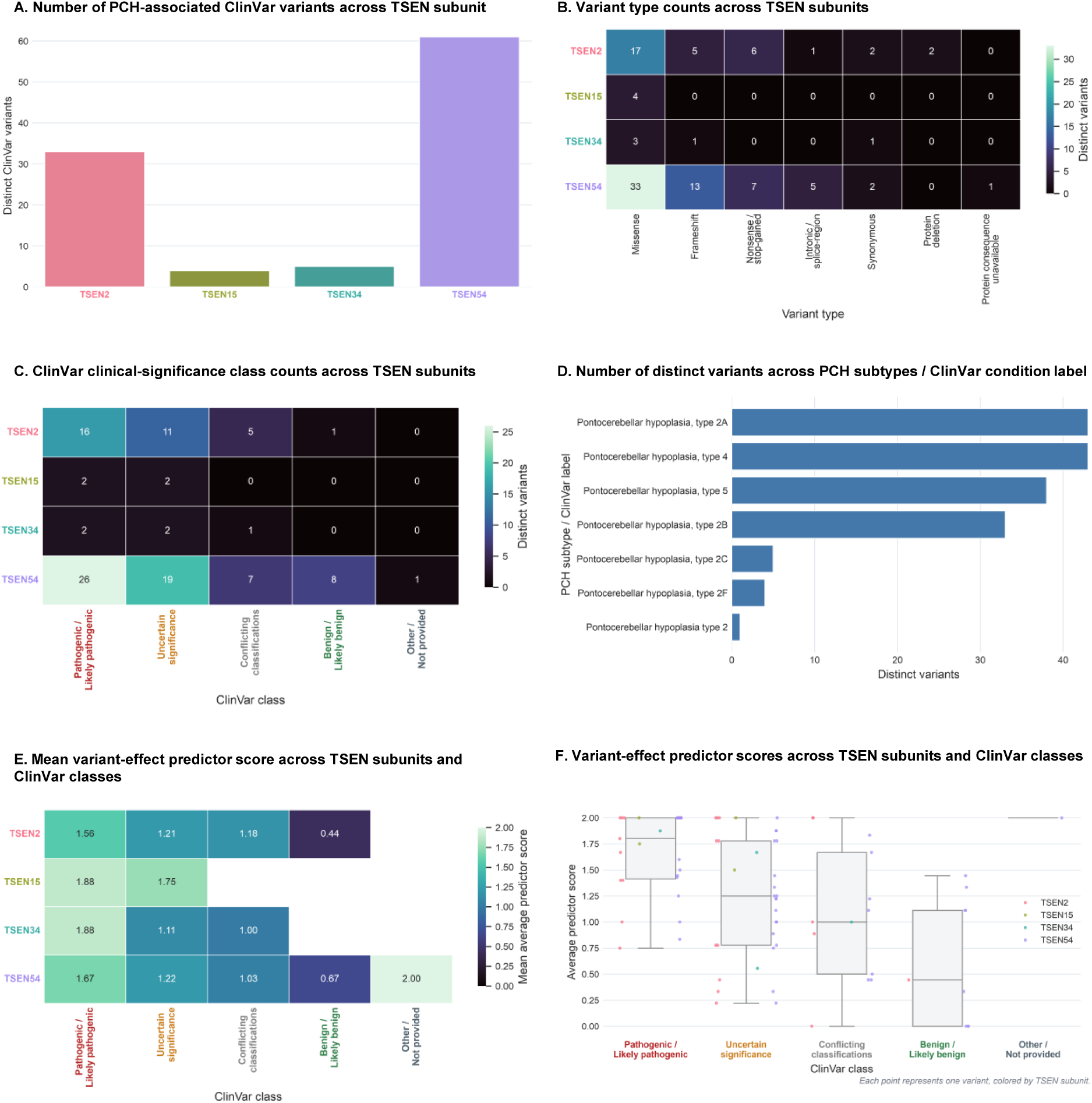
Characterisation of PCH-associated ClinVar variants across TSEN subunits. A. Number of PCH-associated ClinVar variants across TSEN subunit. B. Variant type counts across TSEN subunits. C. ClinVar clinical-significance class counts across TSEN subunits. D. Number of distinct variants across PCH subtypes / ClinVar condition label. E. Mean variant-effect predictor score across TSEN subunits and ClinVar classes. F. Variant-effect predictor scores across TSEN subunits and ClinVar classes. Each point is one variant and is coloured by TSEN subunit; boxes summarize the median and interquartile range within each ClinVar class.

**Figure Expanded View 2.**
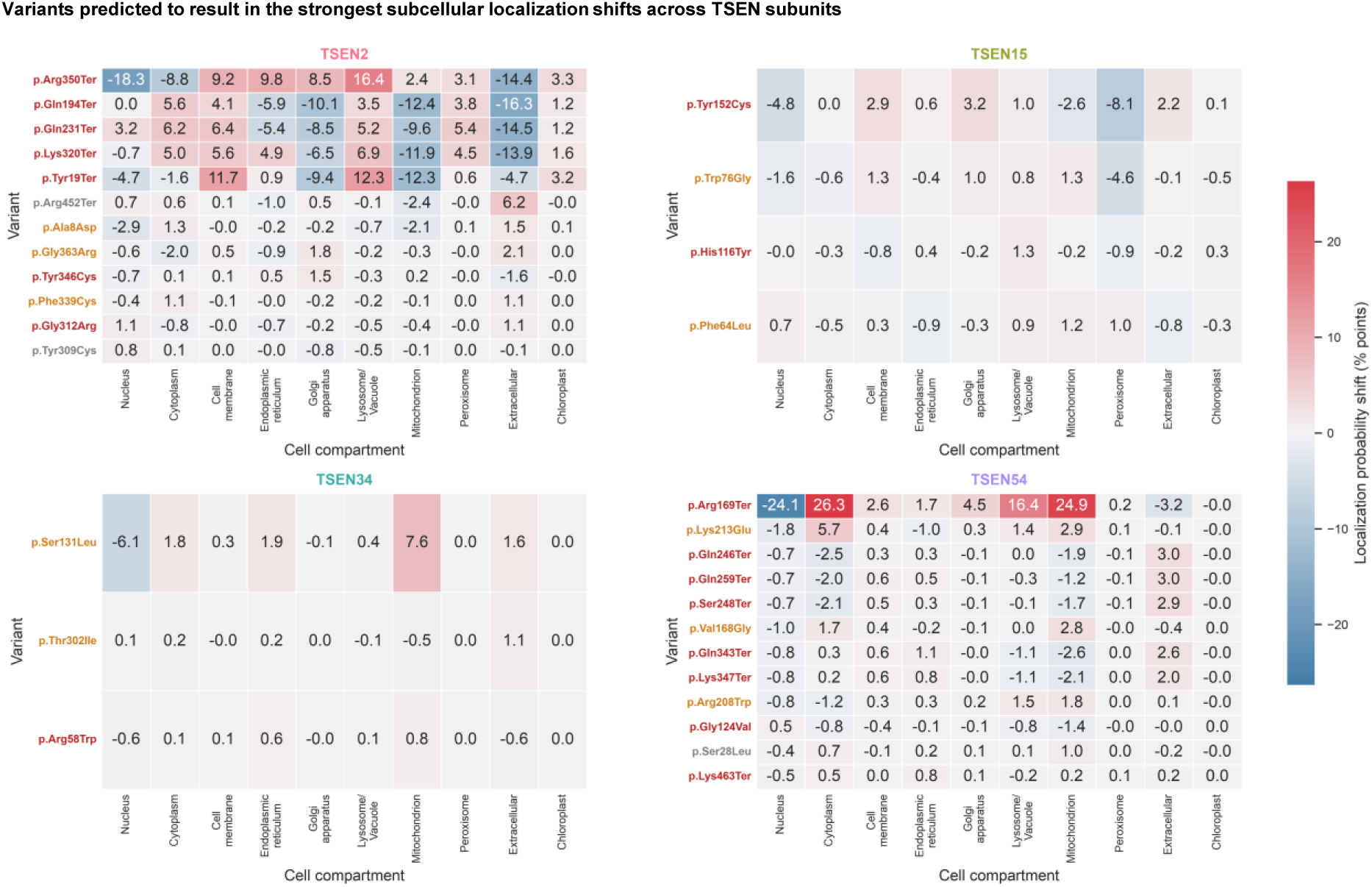
Variants predicted to result in the strongest subcellular localization shifts across TSEN subunits. Localization shifts were calculated as mutant minus wild-type compartment-probability, as predicted with DeepLoc2.1, in percentage points. Variant labels are colored by ClinVar class.

**Figure Expanded View 3.**
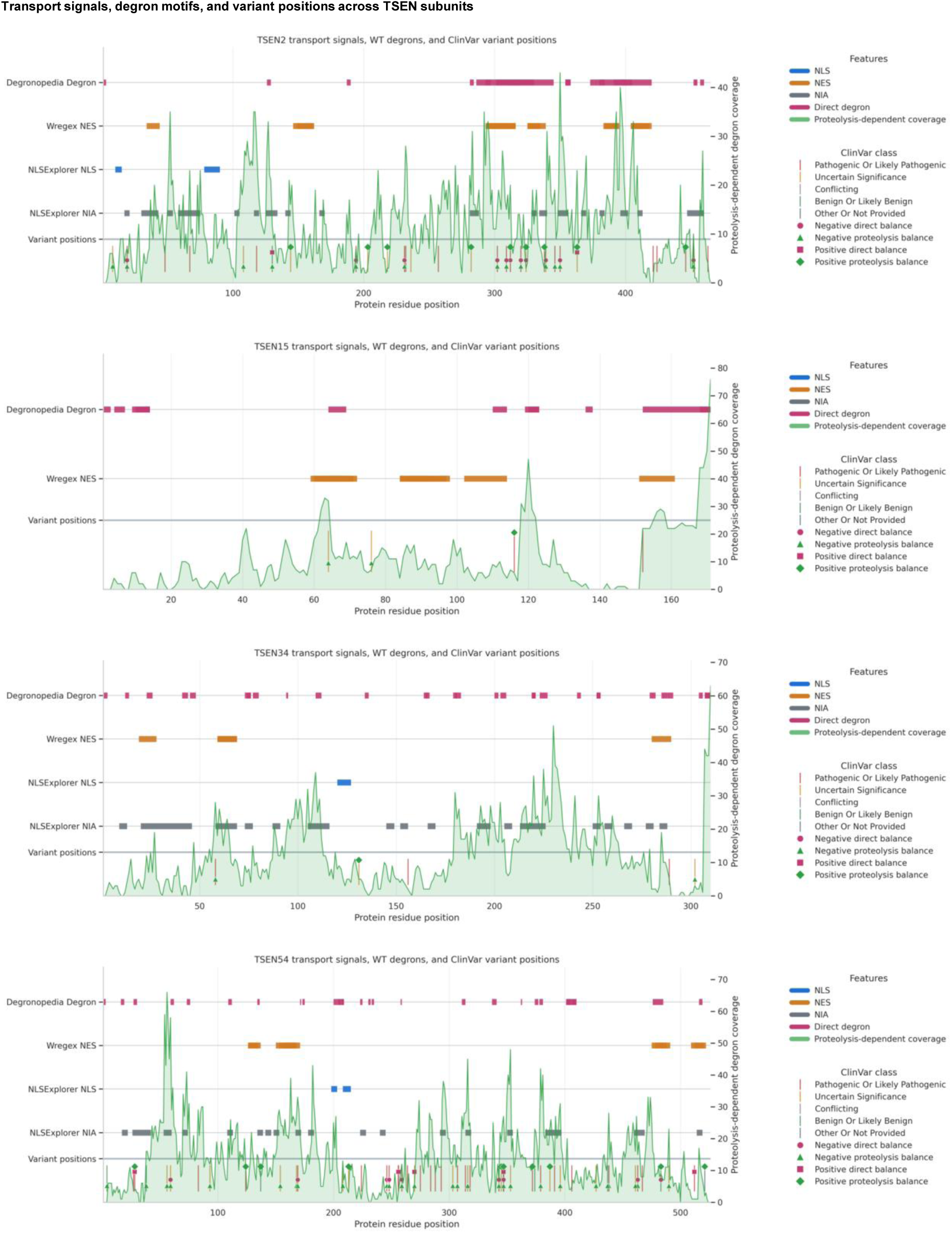
Transport signals, degron motifs, and variant positions across TSEN subunits. Each panel shows predicted transport-signal regions — nuclear localization signals (NLS, from NLStradamus and NLSExplorer), nuclear export signals (NES, from Wregex), and NLSExplorer’s “potential Nuclear Import/Important Area” calls (NIA) — wild-type degron annotations from DEGRONOPEDIA, a proteolysis-dependent degron coverage track (the count of predicted proteolysis-dependent degron regions overlapping each residue), and ClinVar variant positions, for one TSEN subunit. Variant tick marks are further annotated with whether that mutation is predicted to gain or lose a direct or proteolysis-dependent degron.

**Figure Expanded View 4.**
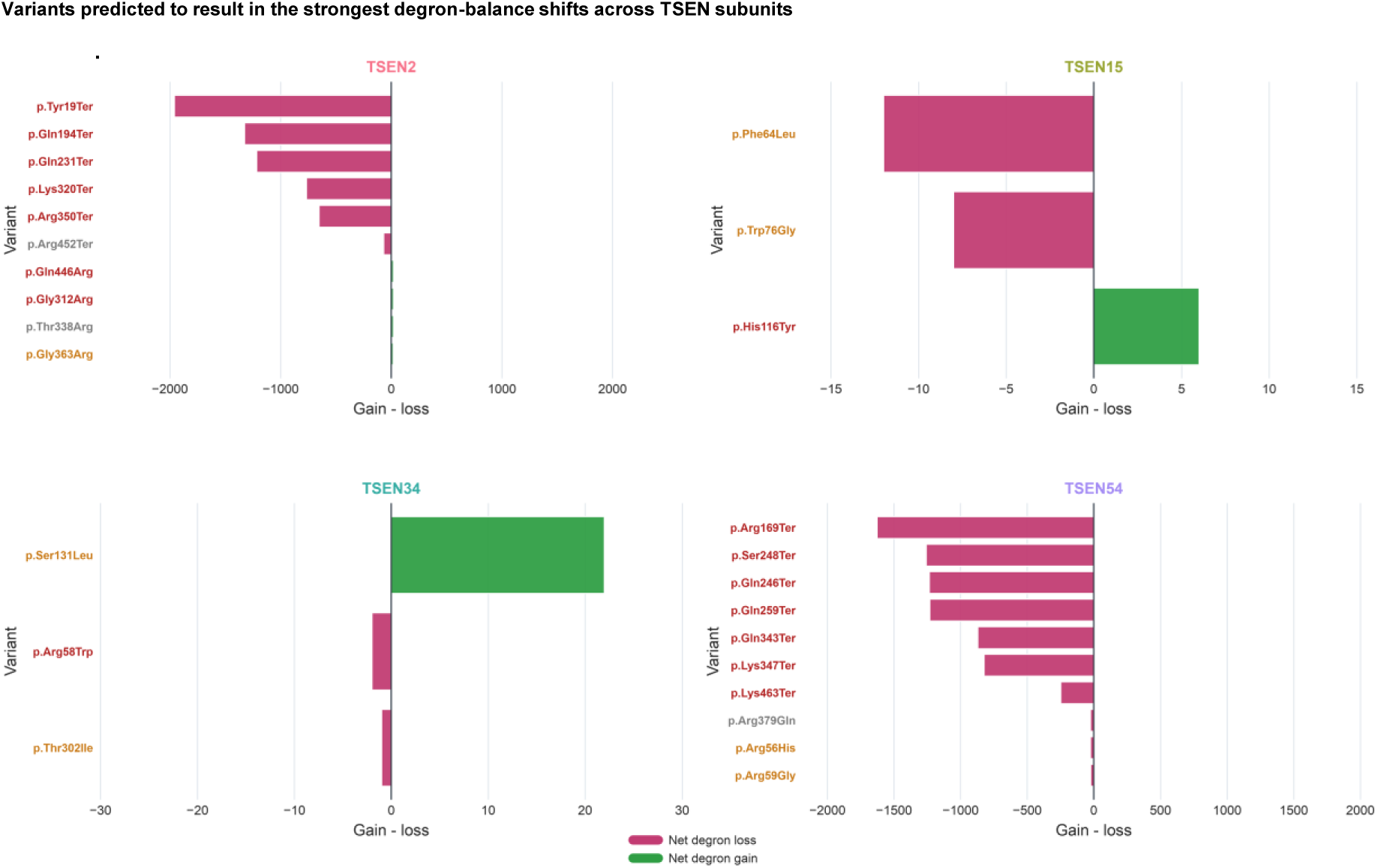
Variants predicted to result in the strongest degron-balance shifts across TSEN subunits. Degron balance shifts are quantified as the difference between the number of gained and lost degrons (direct and proteolysis-dependent). Variant labels are colored by ClinVar class.

**Figure Expanded View 5.**
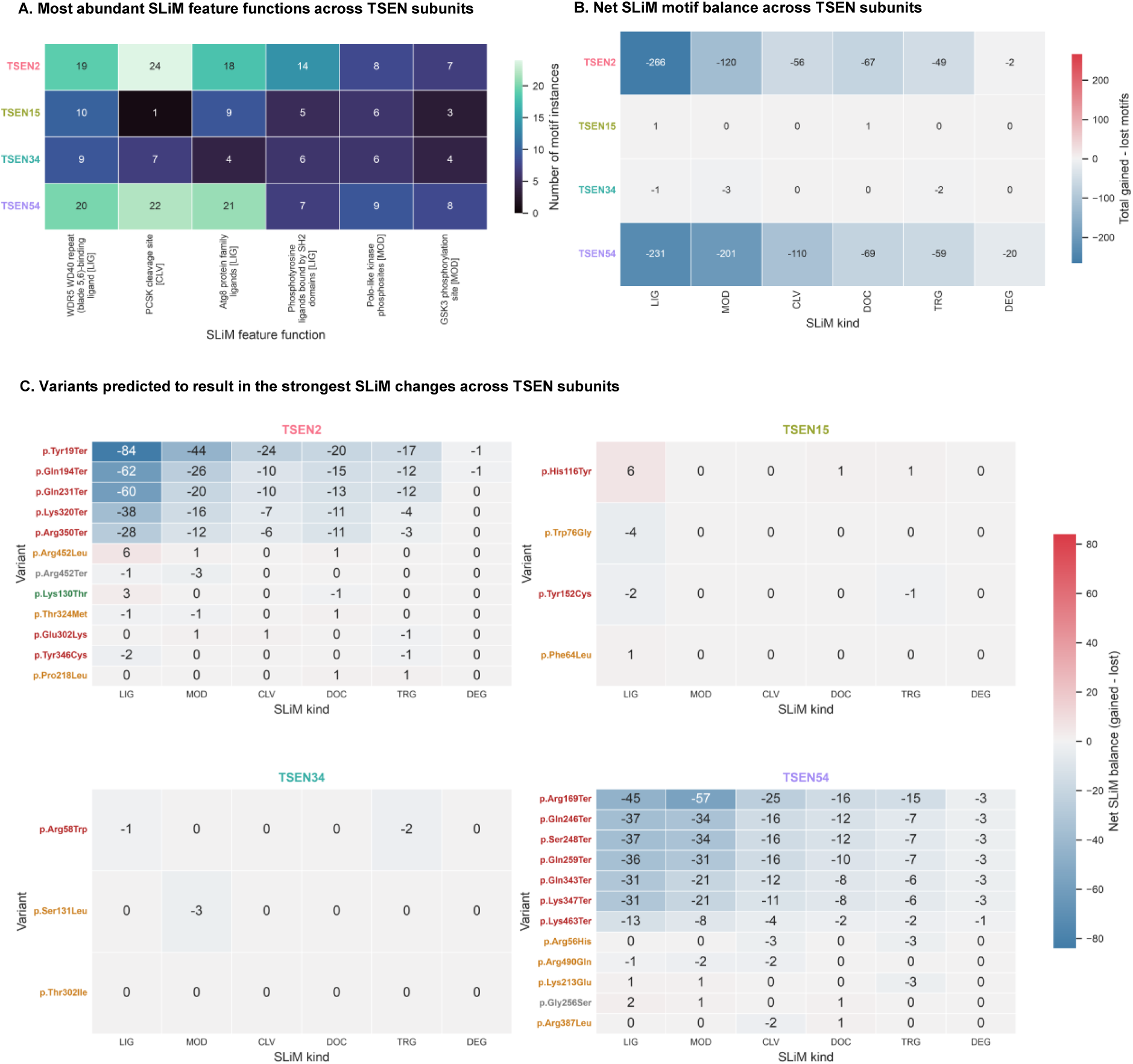
Variant-induced changes in short linear motifs (SLiMs) across TSEN subunits. A. Top 6 most abundant SLiM feature functions across TSEN subunits. B. Net SLiM motif balance across TSEN subunits. Positive values indicate more gained than lost motifs across variants; negative values indicate net motif loss. C. Variants predicted to result in the strongest SLiM changes across TSEN subunits. Positive values represent motif gain, negative values represent motif loss, and variant labels are colored by ClinVar class. *Data information:* In (A-C), the following abbreviations are used for the different classes of SLiMs (ELM motif classes): LIG, ligand-binding sites; MOD, post-translational modification sites; CLV, cleavage sites; DOC, docking sites; TRG, targeting signals (motifs directing subcellular localization or trafficking, e.g. sorting and retention signals); and DEG, degradation sites.

**Figure Expanded View 6.**
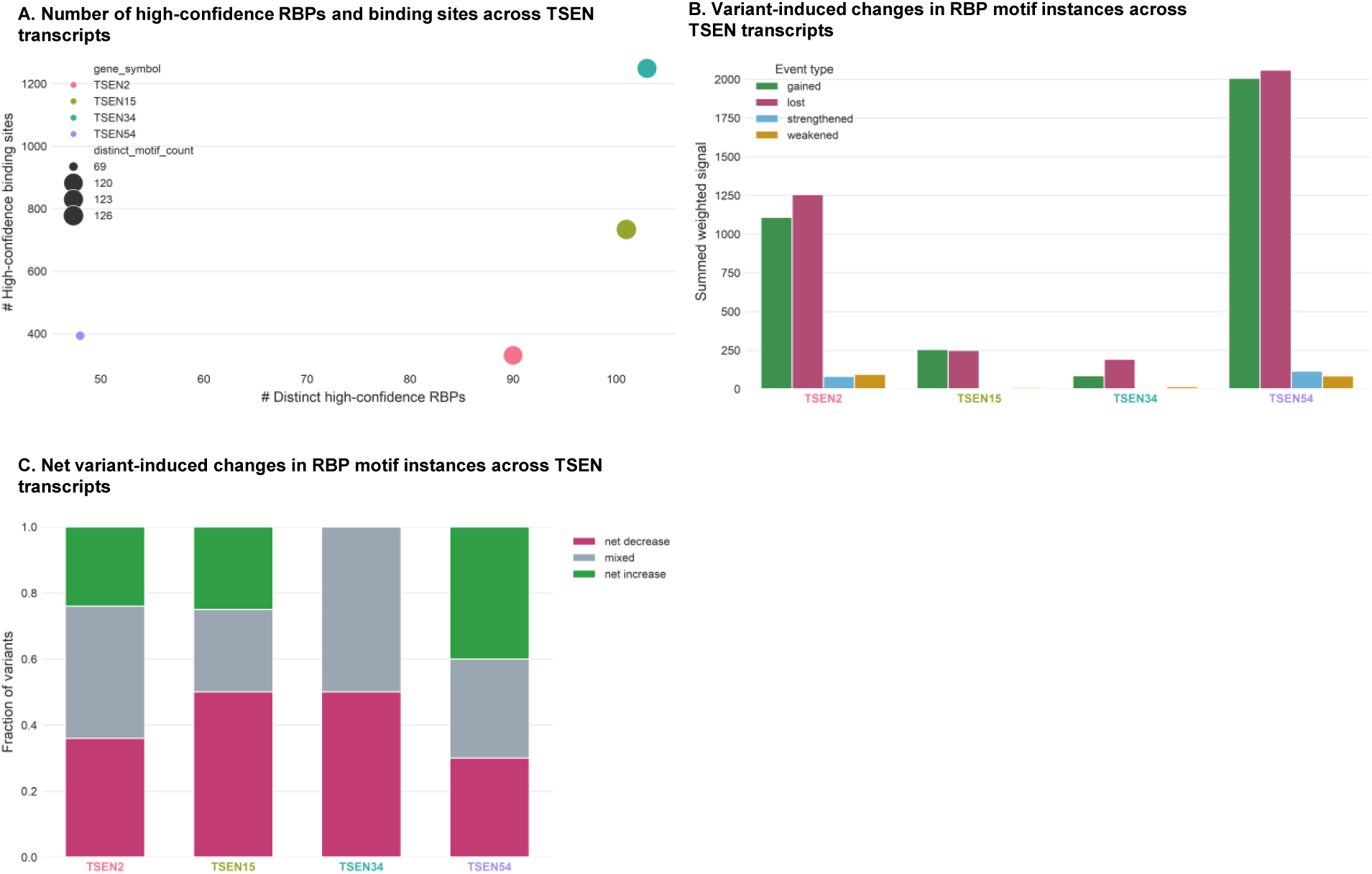
Variant-induced changes in RNA-binding protein (RBP) motifs across TSEN transcripts. A. Number of high-confidence RBPs and binding sites across TSEN transcripts. Each point represents a transcript without variant, colored by gene and positioned by the number of distinct high-confidence RBPs and total high-confidence binding sites; point size reflects motif diversity (the number of distinct RBP:motif label pairs). High-confidence binding sites were filtered using a Z-score > 3 threshold. B. Variant-induced changes in RBP motif instances across TSEN transcripts. Variant-induced changes are separated into gained, lost, strengthened, and weakened binding-site evidence. Unlike panel A, panels B-C use RBPmap’s default reporting threshold (P-value < 0.05) and not the stricter Z-score > 3 cut-off used to define “high-confidence” sites. C. Net variant-induced changes in RBP motif instances across TSEN transcripts.

**Figure Expanded View 7.**
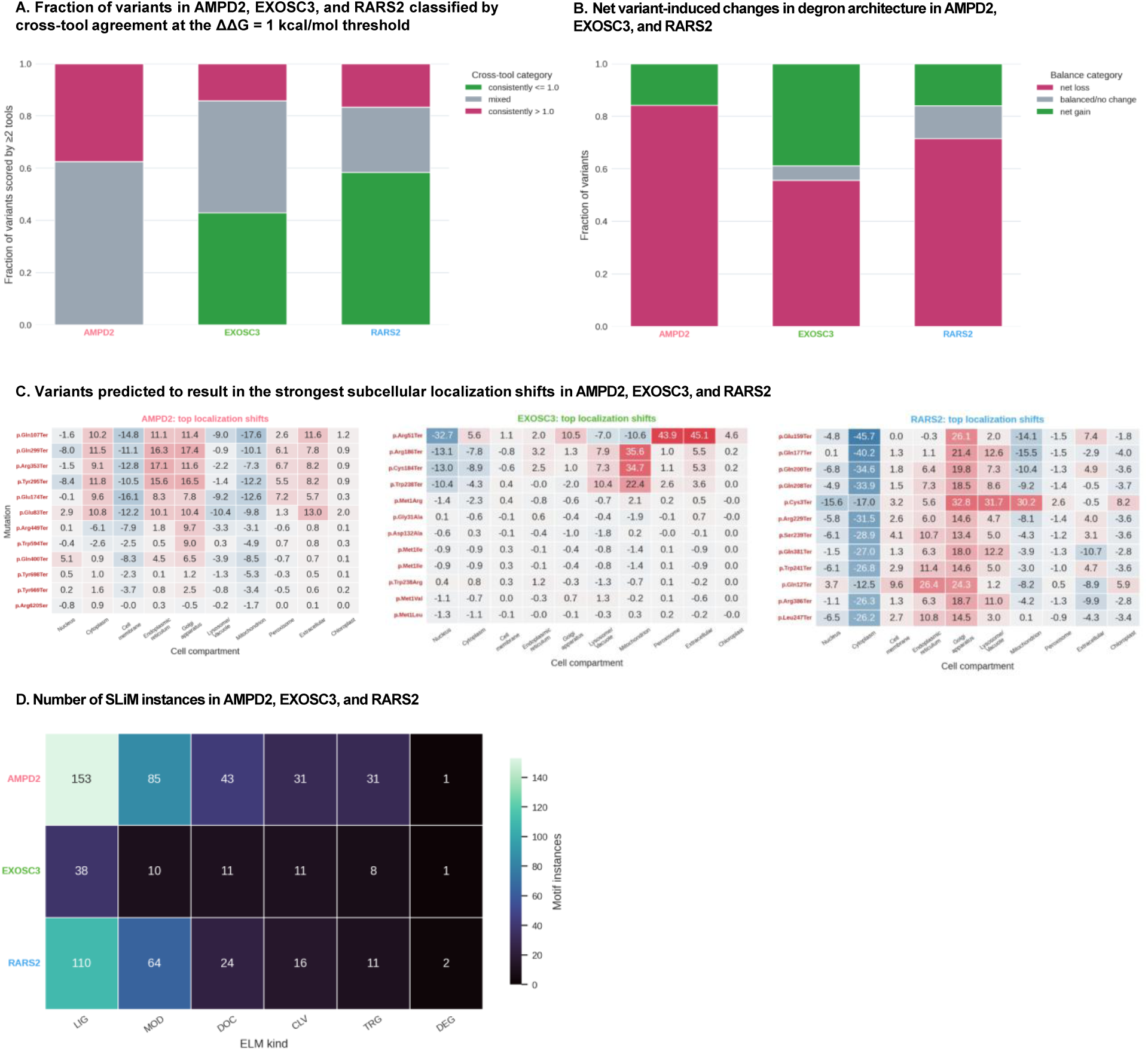
Predicted variant-induced changes in protein stability, subcellular localisation, degron architecture, and short linear motifs in AMPD2, EXOSC3, and RARS2. A. Fraction of variants in AMPD2, EXOSC3, and RARS2 classified by cross-tool agreement at the ΔΔG = 1 kcal/mol. B. Net variant-induced changes in degron architecture in AMPD2, EXOSC3, and RARS2. Each stacked bar depicts the fraction of variants per protein classified as net degron loss, balanced/no change (gained degrons exactly equal lost degrons), or net degron gain, based on the difference between mutant and wild-type degron counts, as predicted with Degronopedia. C. C. Variants predicted to result in the strongest subcellular localization shifts in AMPD2, EXOSC3, and RARS2. Localization shifts were calculated as mutant minus wild-type compartment-probability, as predicted with DeepLoc2.1, in percentage points. Variant labels are colored by ClinVar class. D. Number of SLiM instances in AMPD2, EXOSC3, and RARS2, grouped by motif class.

**Figure Expanded View 8.**
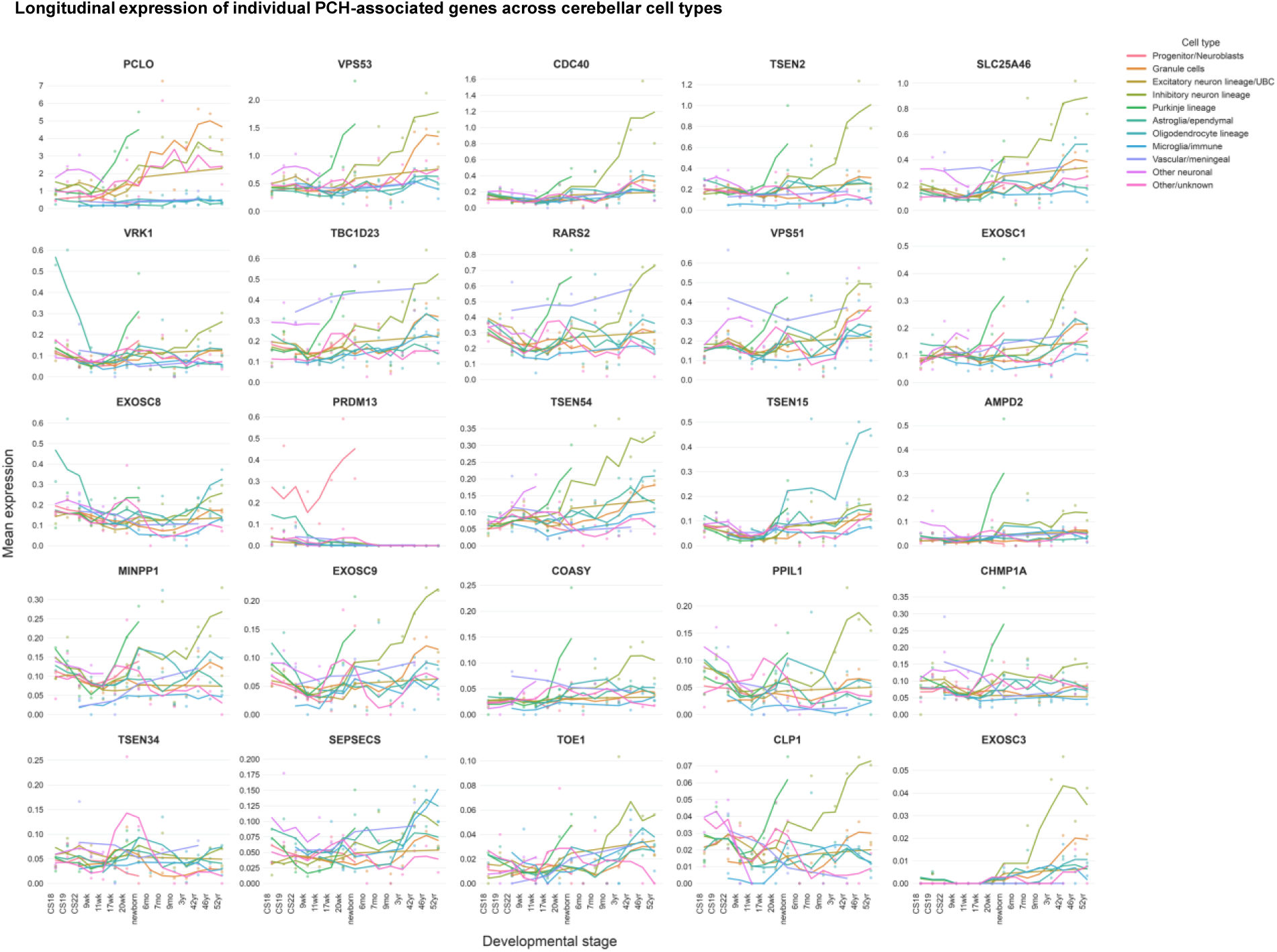
Longitudinal expression of individual PCH-associated genes across cerebellar cell types in the human cerebellar dataset from Sepp *et al*, 2024. Each panel represents one gene and each colored curve represents one harmonized cerebellar cell type (Table 9). Points show observed stage-level pseudobulk mean expression, while lines show a centered three-stage rolling average.

**Figure Expanded View 9.**
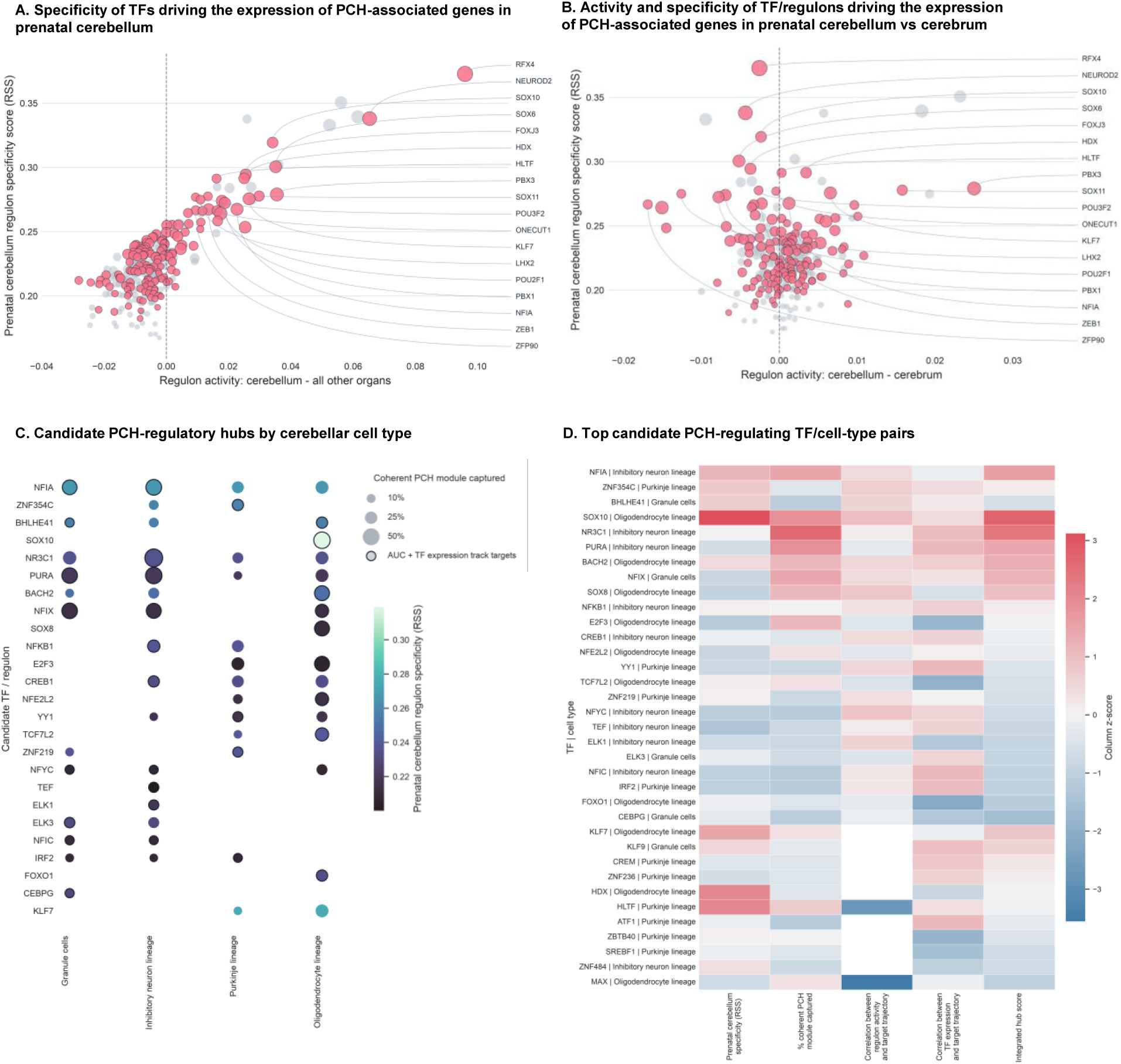
Candidate transcription factors regulating coherent modules of PCH-associated genes in a cell-type-specific manner across cerebellar development. A. Specificity of TFs driving the expression of PCH-associated genes in prenatal cerebellum, as assessed with SCENIC and a human prenatal transcriptomic atlas (Cao *et al*, 2020). The x-axis shows mean regulon AUCell activity in the cerebellum minus the average of the other organs’ mean AUCell activities; the y-axis shows the cerebellum regulon specificity score. Pink circles identify transcription factors that also emerged as candidate PCH-associated regulatory hubs in a single-nucleus transcriptomic atlas of human cerebellar development (Sepp *et al*, 2024); grey points denote all other regulons. Circle size encodes mean AUCell regulon activity in the cerebellum. Labels mark the highest-ranking PCH-driver TFs by cerebellar RSS and displayed activity contrast. B. As in (A), but the x-axis shows mean regulon AUCell activity in the cerebellum minus the cerebrum specifically, rather than minus the other organs generally. C. Candidate PCH-regulatory hubs by cerebellar cell type, as assessed with SCENIC a single-nucleus transcriptomic atlas of human cerebellar development (Sepp *et al*, 2024). Rows show candidate TFs and columns show harmonized cerebellar cell types (Table 9). Dot size encodes the percentage of the coherent PCH-associated gene module captured by the regulon in that cell type, while color encodes the cerebellum regulon specificity score assessed in a human prenatal transcriptomic atlas (Cao *et al*, 2020). Black outlines mark TF-cell-type pairs where both fixed-regulon AUCell activity and TF mRNA expression are positively correlated (ρ ≥ 0.35) with the captured PCH-target module’s expression over development. D. Top candidate PCH-regulating TF/cell-type pairs. Four layers of evidence were considered: cerebellum regulon specificity (RSS, as assessed with SCENIC and a human prenatal transcriptomic atlas, computed relative to all other organs), percentage of the coherent PCH-associated module captured, correlation between fixed regulon AUCell activity and the captured PCH-target module, and correlation between TF expression and the captured module. The candidate hub score combines the other four columns shown here, plus the statistical significance of the coherent-module overlap test itself (see Methods). Values are column-wise Z-scores.

**Figure Expanded View 10.**
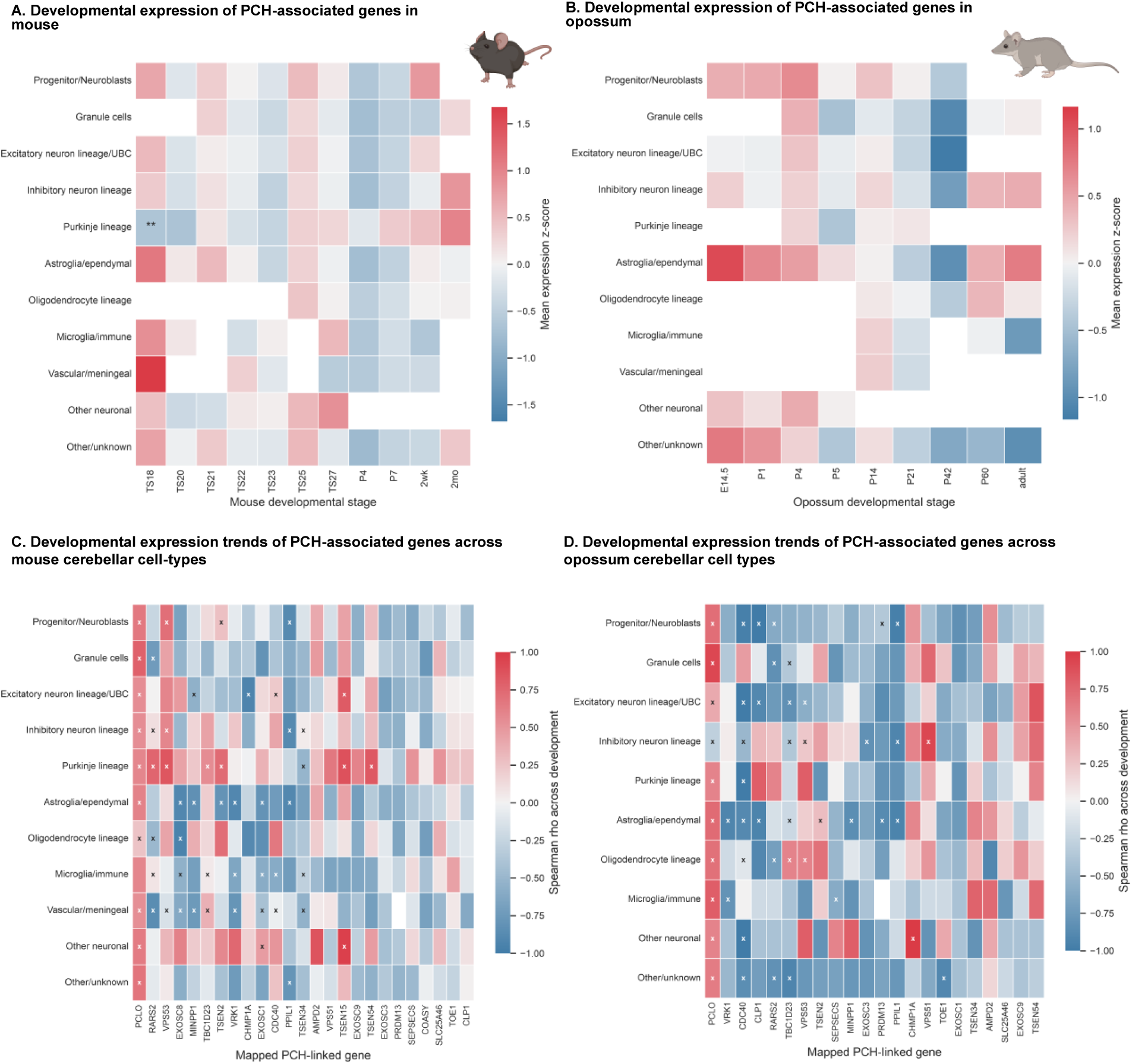
Cross-species comparison of developmental expression trends of PCH-associated genes. A. Developmental expression of PCH-associated genes in mouse cerebellar dataset from Sepp *et al*, 2024. For each of the 11 mouse developmental stages (Theiler stage 18 through 2 months) and each of the 11 harmonized cell types, the plotted value is the mean expression z-score of the PCH-associated genes mapped to their mouse orthologs (all 25 genes mapped). Asterisks indicate cell-type-vs-rest significance within the same developmental stage, assessed by a two-sided Mann-Whitney U test with Benjamini-Hochberg correction across tested cells (* q < 0.05, ** q < 0.01, *** q < 0.001). B. As in (A), but for the opossum cerebellar dataset from Sepp *et al*, 2024, across the 9 opossum developmental stages sampled (E14.5 through adult). PCH-associated genes were mapped to opossum Ensembl orthologs via BioMart human-to-opossum orthology (22 of 25 genes mapped; *COASY*, *EXOSC8*, and *TSEN15* could not be mapped to a feature present in this dataset). No stage-cell-type combination reached q < 0.05 significance, so no asterisks are shown. C. Developmental expression trends of PCH-associated genes in mouse cerebellar dataset from Sepp *et al*, 2024. Each cell is coloured by the Spearman correlation (ρ) between ordered developmental stage and mean gene expression for that gene-cell-type pair. Positive values indicate increasing expression across development; negative values indicate decreasing expression. “x” marks pairs with either a significant monotonic trend (|ρ| ≥ 0.55 and Benjamini–Hochberg-adjusted P-value < 0.1, corrected across all pairs) or, in the absence of such a trend, a stage-peaked pattern (expression range across stages ≥ 85^th^ percentile of all pairs). Unmarked pairs met neither criterion. D. As in (C), but for the opossum cerebellar dataset from Sepp *et al*, 2024, restricted to the 22 PCH-associated genes mappable in this dataset (see panel B).

## Tables

**Table 1. Reference catalogue of pontocerebellar hypoplasia (PCH) subtypes.** Each row corresponds to one OMIM phenotypic-series PS607596 disease entry and reports the PCH subtype, OMIM disease identifier, associated causal gene and respective function description. This table defines the PCH-associated gene set used in downstream single-cell, cross-organ, and regulatory-network analyses.

**Table 2. Variant-level annotation table for all ClinVar PCH-associated variants in TSEN-complex subunits retained in the study.** ClinVar fields report the PCH subtype labels, condition terms, review status, and normalized pathogenicity class used throughout the analyses. Variant-effect columns combine raw or source-scale values from Ensembl VEP/associated annotations, including SIFT, PolyPhen-2, REVEL, AlphaMissense, CADD, phyloP, phastCons, GERP++, and gnomAD allele-frequency information, with project-normalized predictor classes. For the normalized deleteriousness summary, each available predictor call was encoded as benign=0, intermediate/uncertain=1, or deleterious/pathogenic=2, and the average predictor score was calculated as the sum of available encoded calls divided by the number of predictors that produced an interpretable value; missing predictors were excluded from the denominator and reported separately.

**Table 3. Structural-stability table restricted to missense variants TSEN-complex subunits**. The table reports ClinVar/PCH metadata together with per-tool changes in Gibbs free energy or tool-specific destabilization scores from FoldX, mCSM, DynaMut2, and ThermoMPNN where available. Scores are reported in kcal/mol when produced or converted by the corresponding workflow; positive destabilization scores indicate predicted reduction in protein stability, whereas negative values indicate predicted stabilization. Consensus columns summarize the number of methods with numeric output, the methods available for each variant, and the mean, median, and maximum predicted destabilization across scored tools.

**Table 4. Variant-induced changes on protein-localization and degron-architecture across TSEN subunits.** Subcellular localization columns show, for each compartment, the difference in DeepLoc 2.1 probability between the mutant and the reference protein sequence (mutant minus reference); positive values indicate increased predicted localization probability in the mutant sequence and negative values indicate decreased probability. Degron columns summarize direct and proteolysis-dependent degrons annotated with Degronopedia by comparing each mutant sequence with its reference sequence. Total gained and lost degron counts are reported separately, and net degron balance is calculated as gained minus lost motifs.

**Table 5. Variant-induced changes on short linear motif (SLiM) across TSEN subunits.** SLiM instances in each mutant protein sequence were compared with those in the corresponding reference sequence. Instances present only in the mutant were classified as gained, and those present only in the reference as lost. For each variant, the table reports the number of gained and lost SLiM instances and semicolon-delimited motif annotations including motif/function name, SLiM kind, and amino-acid position or interval in the relevant sequence context.

**Table 6.** Variant-induced changes on RNA-binding protein (RBP) motifs across TSEN subunit transcripts. RBP motif gains and losses were identified by comparing the motif instances predicted for each mutant transcript sequence with those from the corresponding reference transcript. Weighted signal values combine motif statistical support and Z-score evidence as determined with RBPmap (see Methods); positive signed signal corresponds to motif gain or strengthened binding signal, whereas negative signed signal corresponds to motif loss or weakened signal. Reported positions are RNA/transcript coordinates.

**Table 7. Integrated variant-induced changes on protein structure and function across TSEN-complex subunits.** Each row corresponds to one ClinVar PCH-associated TSEN variant, including missense, frameshift, nonsense/stop-gained, splice-region/intronic, synonymous, deletion, and variants lacking confident protein-consequence annotation. The table joins ClinVar/PCH metadata, variant-effect prediction scores, DeepLoc2.1 localization deltas, degron balance, SLiM changes, and RBPmap motif changes for all variants; structural-stability fields are populated only for variants meeting the single missense substitution criteria required by FoldX, mCSM, DynaMut2, and ThermoMPNN.

**Table 8. Integrated variant-induced changes on protein structure and function in RARS2, EXOSC3, and AMPD2.** The table follows the same schema as the TSEN all-variant integrated table (Table 7), combining ClinVar/PCH metadata, variant-effect predictions, structural-stability outputs where applicable, DeepLoc2.1 localization deltas, degron balance, SLiM changes, and RBPmap motif changes.

**Table 9.** Mapping between the original and harmonized cell-type labels used in this study when analyzing the single-nucleus transcriptomic atlas of human cerebellar development (Sepp *et al*, 2024).

**Table 10. Candidate transcription factor (TF)-cell type regulatory hubs behind PCH pathology.** Rows correspond to TF-cell type pairs, ranked by an integrated candidate-hub score that combines: the TF/regulon’s specificity score in the prenatal cerebellum (“Cerebellum RSS”, computed relative to all other organs); the percentage of a coherent module of PCH-associated genes captured by the regulon in that cell type (“Percent coherent PCH module captured”); a combined longitudinal-support term averaging the correlation (r) of regulon activity (AUC)/TF expression with the captured PCH-gene module (“AUC-target trajectory r”/”TF-target trajectory r”); and the statistical significance of the coherent-module overlap test itself. This table also lists the exact number and number of PCH-associated genes that integrate each of the listed TF-cell type regulatory hubs.

